# Charge-trap flash memory cells of the brain

**DOI:** 10.64898/2026.06.29.733154

**Authors:** Philip P. Foster, Raj S. Chhikara, Aladin M. Boriek

**Affiliations:** University of Houston-Clear Lake, Department of Mathematical, Applied, and Physical Sciences, Houston, Texas, USA; Baylor College of Medicine, Department of Internal Medicine, Houston, Texas, USA; ConceP.T, Houston, Texas, USA

## Abstract

Despite extensive study of cellular mechanisms underlying long-term potentiation, no single specific protein or gene has been identified which encodes an individual unit of information, or memory bit. Indeed, the brain engram remains a knowledge gap. Guided by hypothesis-driven statistical analyses of biological data, it suggested two distinct domains of investigation: biological macroscopic measurements and quantum-scale mechanisms. The theory of exclusion led us to cancel one-by-one biologically implausible alternatives. We present evidence supporting quantum tunnelling as a potential mechanism for memory storage within the ubiquitous near-ideal topological insulator located in-between myelin layers, specifically in the paranodal and juxtaparanodal regions, but not at the nodes of Ranvier. Superposition of up to concentric 300 myelin layers, spiraled, and highly compacted wrapping a single axon and each wrap could host hundreds to thousands of niches, as “memory cells”, collectively consisting of a massive array of cells. The disjointed 3D spatial superposition allows storage of charges, nodes not facing from a layer to next. The thickness of a single myelin layer ranges from 7.0 to 20 nm. The dimension scale is approximately the exact dimensions of the charge trap, the tunnel and dielectric also equipping current AI microchips. Stored charges are positive ions, with similar effect whether charges are negative or positive charges creating an electromagnetic field. To “write” data, following an action potential, this voltage applies to the control gates of the myelin layers producing an ionic charge injection. This causes charges to gain energy and “tunnel” through the myelin layer across nodes, via quantum tunneling, and deep into the concentric myelin multilayers. This is creating an insulated trapping of *K*^+^ ions isolated from the system. In a long white matter tract bundle, the near-perfect isolation of millions of axons within compressed myelin wrap-ion channel *K*^+^*/Na*^+^ systems provides quantum coherence and precision of asynchronous firing property. The injected ionic charges (*K*^+^) become physically stuck in “traps” within the myelin layers. The *K*^+^ ions may not move freely, completely trapped after AP ceases. Mirroring a single-bit, single-level-cell, a trapped ionic charge (ions *K*^+^) may represent a “1,” while an empty cell (absence of *K*^+^) represents a “0”. The trial-and-error process, with a Bayesian inference which may have also been the core evolution of the learning human brain. Based on selected mathematical equations, we analyzed the general scheme on how deep learning may be embedded in the brain.

## A. Introduction and background

Understanding how information is encoded continues to elude current research efforts. The vast number of neurons and synapses, combined with dynamic processes such as plasticity, continuous synaptic remodeling, and reinforcement-dependent pruning, creates a severe curse of dimensionality within high-dimensional spaces. This complexity makes it extremely difficult to identify and track fundamental units of information within and across dimensions using a unified approach. Although biological systems operate within the laws of physics, the complexity of high-dimensional biological spaces renders them seemingly resistant to reductionist analysis. Consequently, cross-disciplinary research is required—one that can integrate fragmented yet interpretable “small worlds,” ranging from single-neuron transcriptomics to deep and reinforcement learning, into multilayered networks reminiscent of those in artificial intelligence. Confronted with similarly high-dimensional data and inspired by brain architecture, the AI field has developed methods to address these challenges. Nevertheless, efforts to meaningfully reconcile artificial intelligence with the human brain remain unfruitful. Confronted with vast high-dimensional datasets, AI has achieved simplification through iterative trial-and-error processes, often grounded in Bayesian inference—an approach that may also reflect a fundamental evolutionary principle of human brain learning. Recent advances in artificial intelligence and machine learning (ML) include supervised, self-supervised, and reinforcement learning paradigms. Deep learning (DL) employs multilayered artificial neural networks trained through error-correction mechanisms such as backpropagation and feedforward propagation^1,2.^ Despite these advances, efforts to align AI mechanisms with human brain function have largely been unsuccessful. We initiated this study by examining how machine learning, particularly deep learning, could potentially be embedded in the brain. Consistent with prior investigations, we encountered a persistent obstacle: the lack of empirical evidence supporting the notion that the human brain operates as a digital system, a limitation previously recognized as a critical barrier^3^. We selected the fundamental equations of deep learning which constitutes one of the modalities by which the human brain learns. Importantly, the brain does not compute elementary statistics directly; rather, these equations serve as computational analogues to replicate observed cognitive processes. Nevertheless, subtle interactions among temporal dynamics, biometric constraints, and energy consumption suggested the existence of underlying information vectors at sub-levels. This observation motivated further exploration at the nano- and quantum scales. While investigations at the nanoscale did not yield satisfactory models, analysis of AI-microchip architecture revealed that memory cells constitute the fundamental unit of information storage. Subsequent examination of quantum-level phenomena highlighted mechanisms potentially supporting data retention within these cells.

In the subsequent phase of our study, we implemented two comprehensive inventories: (i) theoretical and experimented models of memory formation, and (ii) inventory of the brain equipment potentially capable of supporting long-term memory storage, commonly referred to as the engram. While the propagation of information via action potential is well characterized, the biological and structural substrates underlying long-term information storage remain largely unknown. Inferences drawn from the absence of experimentally verified AI-microprocessor storage mechanisms may provide insight into long-term potentiation (LTP) and memory formation. Long-term potentiation was extensively studied albeit the brain engram remains a knowledge gap. We performed a series of hypothesis-driven analyses of biological data that yielded highly statistically significant results. However, these comparisons involved fundamentally different measurement scales. Thereby, it became clear that the results suggested two distinct domains of investigation: biological macroscopic measurements and quantum-scale mechanisms. Furthermore, under a purely biologically driven hypothesis, and absence of direct specific quantum observational data, we performed Monte Carlo simulations to assess whether the observed findings could be interpreted without direct measurement of putative quantum mechanical effects. Owing to the fundamental limitations on direct measurement at the quantum scale, as imposed by the Heisenberg uncertainty principle, the subsequent step relied on exclusionary reasoning. This included strategies analogous to Michelson–Morley-type arguments, enabling the systematic elimination of biologically implausible mechanisms. The theory of exclusion led us to cancel one-by-one several unrealistic biological options, suggesting that the explanation resides somewhere else. Collectively, these findings indicated that the underlying solution may lie beyond the scope of conventional biological frameworks. A detailed examination of the brain’s myelin sheath, synaptic plasticity and strength, compared to analogies with stable charge-trap flash memory and the fast accessibility of dynamic RAM (specifically the 1T1C fundamental memory cell), yields further insight.

Clearly, investigating machine learning, data writing and memory processing in the brain led us to the foundations for a binary process. All of which guided us to inspect the brain at the quantum scale and achieve an inventory of its structures potentially supporting data writing and storage. Hereby, we started the investigation by looking at the available engineered models from AI computing. Accordingly, our investigation was initiated through an analysis of existing engineered models derived from artificial intelligence computing. It is evident that examining AI machine learning, information encoding, memory processing and potential parallels in the brain directed our attention to an underlying binary framework. This, in turn, motivated us to investigate the brain at the quantum scale to catalogue structures which might support information encoding and storage, and we completed a full inventory.

This work emphasizes the core mathematical principles that support these exclusions and provides hints on one model of brain systematization. Thereafter, we investigated the brain cell machinery, neurons, oligodendria, microglia, glia (astrocytes), and did an inventory of the equipment which could support the engram. Our analysis was derived from quantum physics principles applied in AI microchips. Envisioning the digitalization is fully achievable through this binary framework (1 or 0) in the physiological brain charge trap flash memory cells. This exhaustive analysis revealed an amazing engineering design, favoring a packed, multi-function circuitry, to optimize the volume, weight, energy, efficiency, rapidity, precision, storage, and same conductive input circuitry reusable in memory cells. Addressing the complex multidisciplinary issue of brain action potential demands careful categorization and ***in-depth analysis*** of a wide range of interconnected elements, *including those whose relationships are not immediately apparent*.

The organization of this paper follows a logical rationale and is structured as follows. **A. Introduction & background**. Section A-I The fundamentals of artificial intelligence (AI), deep learning, and their underlying algorithms, highlighting historical parallels with brain function^1^. It also discusses the evolution of today’s AI toward artificial general intelligence (AGI) and world models. This section establishes the conceptual foundation of a potential “digital brain,” which is further explored in subsequent sections. Section A-II examines a human-engineered model of information storage: *charge-trap flash* (CTF) memory technology, widely used in modern electronic devices. Subsection A-II-1 reviews the principles of quantum tunnelling underlying CTF memory operation. Subsection A-II-2 proposes a novel interpretation of transmembrane ion *transversal transfer* across axons, whereby *K*^+^ and *Na*^+^ ion-channel dynamics are considered from the perspective of *quantum tunnelling*. This ATP-driven transport mechanism moves ions against their electrochemical gradients while minimizing overall energy-processing demands and potentially contributing to the preservation of *quantum coherence* (Fig. 3. and C. Discussion). Subsection A-II-3 introduces a complementary mechanism involving the *longitudinal propagation* of electromagnetic potentials along axons, where node-to-node transfer of charged particles is considered within a quantum-tunnelling framework relevant to memory storage. Section A-III investigates the *multidimensional* organization of the brain and its role in information transfer. **B. Results**. Subsection B-I-2, excludes implausible candidates for the physical substrate of memory. Subsection B-I-3, identifies potential brain machinery for the *engram*. Section B-II, proposes a potential mechanism underlying the *engram*. **C. Discussion**. Discussion of the overall quantum coherence of the isolated system of myelin wrap-ion channel *K*^+^*/Na*^+^. Particular attention is given to how the layered architecture of neural tissue may provide isolation from environmental disturbances and thereby mitigate *macroscale quantum decoherence*. Subsection C-I-5 is about unique particularity of node, paranode and juxtaparanode in quantum mechanics. Subsection C-II is about the special case of Alzheimer’s disease providing further evidence. C-III. **Conclusion**, the brain represents a highly optimized architecture for memory, information processing, and adaptation of the biped-human in terrestrial one-gravity. **D. Methods**. Focusing on recent biometrics data, distinct velocities of action potential and electromagnetic-driven phase transitions. Subsection D-III, computational statistical modeling using Monte Carlo simulations. **E. Appendix**. *R* Program.

### A-I. Hints that some machine learning-analogous processes may be embedded in the human brain

#### 1. Some brain processes may exhibit deep learning analogous principles

In AI computing, the deep reinforcement learning procedure, and back-propagation is a multilayer of neural networks. The definition of “artificial neuron” and “artificial neural networks” were designed while facing the “wall” of high dimension data, designers did not know how to write a working program. At the time, converting pixel intensity values of an image into a string of words that described the image was deemed impossible. AI seems a phenomenal tool to read and treat information at a non-human pace. We will evaluate how AI may be inspiring to unfold some aspects of the brain operations. Indeed, a mismatch seems to exist between AI computing and operations of anatomical structures of the brain. The generalization theory, the stochastic descent gradient, or cascade of classifiers, multilayer networks, “high-levels”, “mid-levels”, “low-levels” in deep-learning may be imagined in the brain based on the physiology. Physiological frontiers/classifiers may not strictly fit AI’s classifiers and weighing (see Fig. 4b). The human brain anatomy and physiology are the product of a long evolution to optimize the adaptation to the environmental world. The human brain is a model of ingenuity, miniaturization and energy saving. Reproducing human intelligence with a human brain-based machine model rapidly evolved from neural networks and back-propagation^1^, to various machine learning, ML, techniques, such as supervised learning, SL, reinforcement learning, RL, self-supervised learning SSL, feed-forward propagation, and generative prediction^1-5^. Experimental ML relies on Bayesian logic. Made it simple, this can be stated as a gigantic super-fast trial-and-error process, with 1 or 0 outcome, or plausible outcomes. Prior to the blooming of AI, the stochastic aspect of AI was highly disconcerting for programmers writing symbolic/mathematical source code in C++, Mathematica®, or Java and providing specific instructions to the computer. Each algorithm or mathematical symbolic of heuristic models of the human body representing each physiological process was created under strict human guidance. In contrast, AI by trial-and-error process freely solves the question using limited source code in Python or PyTorch or others. We will describe some mathematical aspects of ML in more detail, and we will attempt to include ideas about elemental vectors of information. Although every conceptual brain structure differs from a computer, some framework shows a potential for integrating topological DL with neuroscience^6,7.^

Structurally, biological neural networks and artificial neural networks are differing albeit sharing common features in the macro-dimension MAN, of the multidimensional network architecture shown in Figure 4b. The human brain evolution departs significantly with abilities not relying on speed and treatment power of high dimensional data. In the next section we will focus on fascinating, albeit unexplained features of the brain, as seen on imaging (fMRI). Those features seem to invoke alternate, yet unknown paths for treating information. A few years ago, there was an attempt to use AI-based droids or robots in the US Space Program to achieve or replicate specific human tasks in various outer space extravehicular activity (EVA) missions. Robots-droids were inherently more resistant to the absence of gravity and space void, albeit AI was unsatisfactory in unpredicted situations. All of which led to disappointing results. The human brain reacts in a more appropriate and adapted way to unforeseen threats. Hereby, substituting robots for humans in the Space Program or all aspects of life is yet a challenge. This is known as the Moravec’s paradox^8-11^. Tasks reputed simple for humans often prove difficult for machines, and vice-versa^8-11^. Chess, Go game, daily country’s GDP computations, text generation, next sentence-word prediction in text, translation into over 200 languages and dialects, are no brainer for AI. While an auditory Turing test comprising 917 challenges across seven categories: overlapping speech, speech in noise, temporal distortion, spatial audio, coffee-shop noise, phone distortion, and perceptual illusions using the state-of-the-art audio models (GPT-4’s audio capabilities and OpenAI’s Whisper) returns a failure rate exceeding 93%, with even the best-performing model achieving only 6.9% accuracy on tasks which humans solved at 52% (7.5 times higher success)^11^. AI may sometimes go completely “hallucinating”. There is still a gap between AI and the operating human brain, i.e. world model aka artificial general intelligence. What was remarkable in the rapid development of AI in a couple of decades is the starting point^1^. Oftentimes, inventions and discoveries start with a technological breakthrough. Occasionally, the inception begins with a genius theoretical concep^t1,12-15^ and, decades later, technology/R&D proving it right, augments the initial concept with practical applications, such as large language model, LLM, the foundation of ChatGPT.

Humans learn by vision, audition, smell, touch, emotion, and construct a vision of the world based on feeling the elaborated balance system on two lower limbs, speed, in three dimensions under an impression of elapsing time much different from a simple time-measurement. As always in exploring unknown territory, we must be conscious that we may be forgetting many aspects of our brain, which will be unveiled much later in time. Such a caveat must be made to keep our minds open to more options than those widely advertised about AI today. We will first explore positively how much AI, machine learning and deep learning bring to our understanding of how the brain works. An evidence-based approach leaves machine learning and deep learning as key players in the treatment of information in the human brain. This evidence-based approach relies on anatomy, structure and function, brain imaging, epigenetic priming of memory, DNA double-strand breaks and other brain specifics. Another potential positive hint is based on an intriguing disparity between timing, biometrics, and energy expenditure is raising questions for yet unknown potential mechanisms.

#### 2. Some trial-and-error process in the brain

In AI, a trial-and-error process is postulated to enable the computer recognizing and learning shapes or letters in the environment. This is also a simple modality for training animals to learning tasks and acquiring new skills, like feeding rodents in the lab. Hereby, we are focusing on the fundamental step of machine learning. An appropriate statistical model for the occurrence of an event or the likelihood for a computer to identify a shape or letter is often assumed to be logistic^16-19^. The logistic regression of the sum of trial-and-errors, each one providing a binary outcome (1 or 0), is a sigmoid curve. The probability *P(y)* to correctly identify the shape, is defined by the logistic regression equation, namely

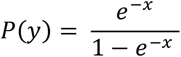

The logistic probability distribution function is widely used in ML^1-5,20,21^. Facing high-dimensional data, this is also the procedure which scientists used in a similar trial-and-error process in the NASA space program^16^. This was a customized-whole-body-procedure to protect the astronaut brain in the NASA space program, while performing extravehicular activities, EVA’s, in outer space^16,22-26^. The maximum likelihood was a preferred technique to estimate unknown parameters in a model so that the probability function is maximized^4,16,20,21,27^. Hereby, the likelihood of identifying the correct letter is optimal. Given a set of observed values (*x*_1_, *…, x*_*n*_) defining the letter, the likelihood function *L* is defined by the product of the probability densities for the letters’ outcomes

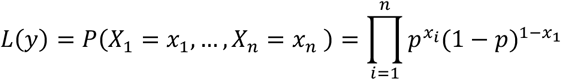

where (*X*_1_, …, *X*_*n*_) are the outcomes, or observed values, by the computer^16^.

It is customary and easier to use the natural logarithm of the likelihood function^16^

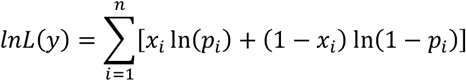

The log likelihood function, *lnL*(*y*), is always negative, since *ln*(*p*_*i*_) and *ln*(1 − *p*_*i*_) are negative as 0 < *p*_*i*_ < 1.

The maximum likelihood estimates of the parameters that maximize the *lnL* function which also minimize the shape/letters reading errors by the machine can be obtained by solving the following equation^16^

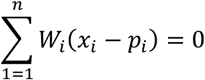

*W*_*i*_ are weight vectors to be adjusted by a trial-and-error process. The procedure is called stochastic gradient descent (SGD). Input vectors are shown for a few examples, with the outcomes and errors obtained, computing the average gradient for those examples, and adjusting the weights correspondingly^2,28^. The process is iterated for several narrow sets of examples from the training set until the mean of the function ceases decreasing. The real world is most often modelled on non-linear analysis of shapes, letters, deduction of the next sentence word, more complex patterns or else. In deep learning, there is an interface of non-linear functions, like the sigmoid, or other non-linear functions, such as the rectified linear unit, ReLU^2,28,29^. The rectified linear unit, ReLU function, *f*(*x*) = *max*(0, *x*), can also be expressed as

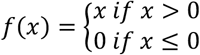

ReLU learns much faster in networks with multiple layers, allowing training of a deep supervised network without unsupervised pre-training^27^.

The interface is made of hidden network layers returning a probability score of reading correctly (0 < *p*_*i*_ < 1), expressed by sigmoid shapes. The sigmoid is a probability “score”. This process is repeated until the correct and/or acceptable answer is provided by the machine. Children are mirroring which facial reaction the adult will reflect. Children instinctively develop their brain through a trial-and-error process^30^. The current architecture of AI is constantly developing and improving from day to day. The most common form of machine learning’s architecture to minimize the error function is supervised learning^2,31-34.^ The rectified linear unit activation, ReLU, in artificial neural networks seems to produce more brain-like and modular neural responses^7^. Repeating this deep learning process through a cascade of classifiers (SGD) ranging from low-level, mid-level, and high-level features, minimizes the error function. What applies effortlessly to the human brain is not easily executed by a computer. This is solved by the convolutional neural network, CNN, analyzes a series of pixel blocks, in which filters operate the pattern recognition. The CNN filter splits the image into modules of much smaller scale. Hereby, the inputs are only a few pixels with the primary colors, RGB (red, green, blue). The computer creates a feature map. After CNNs’ filters reduce the feature map spatial dimension, pooling units compute the maximum units in one or more feature maps. Adjacent pooling units take input by shifting patches. Stages of convolution, non-linearity and pooling are stacked, followed by additional convolutional and fully-connected layers^2^.

#### 3 First stage, recognition of the object

First reading (Fig. 1) flows bottom up. It is a simple trial and error process, progressing by stages, through “layers”. Let postulate the differential *dx* is an independent variable, hereby, applying the chain rule to one step across two layers, L_1_ and L_2_, for two intermediate and independent variables. Let *f* be a non-linear function such as log logistic function (sigmoid) or rectified linear unit, ReLU. Let *f*: ℝ^2^ → ℝ and *g*: ℝ^2^ → ℝ^2^ be differentiable. Write *f* as a function of the variables *u* and *v* and write *g*(*x, y*) = *f*[*u*(*x, y*), *v*(*x, y*)]. Then

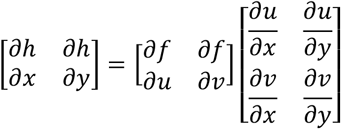

**Fig. 1.**
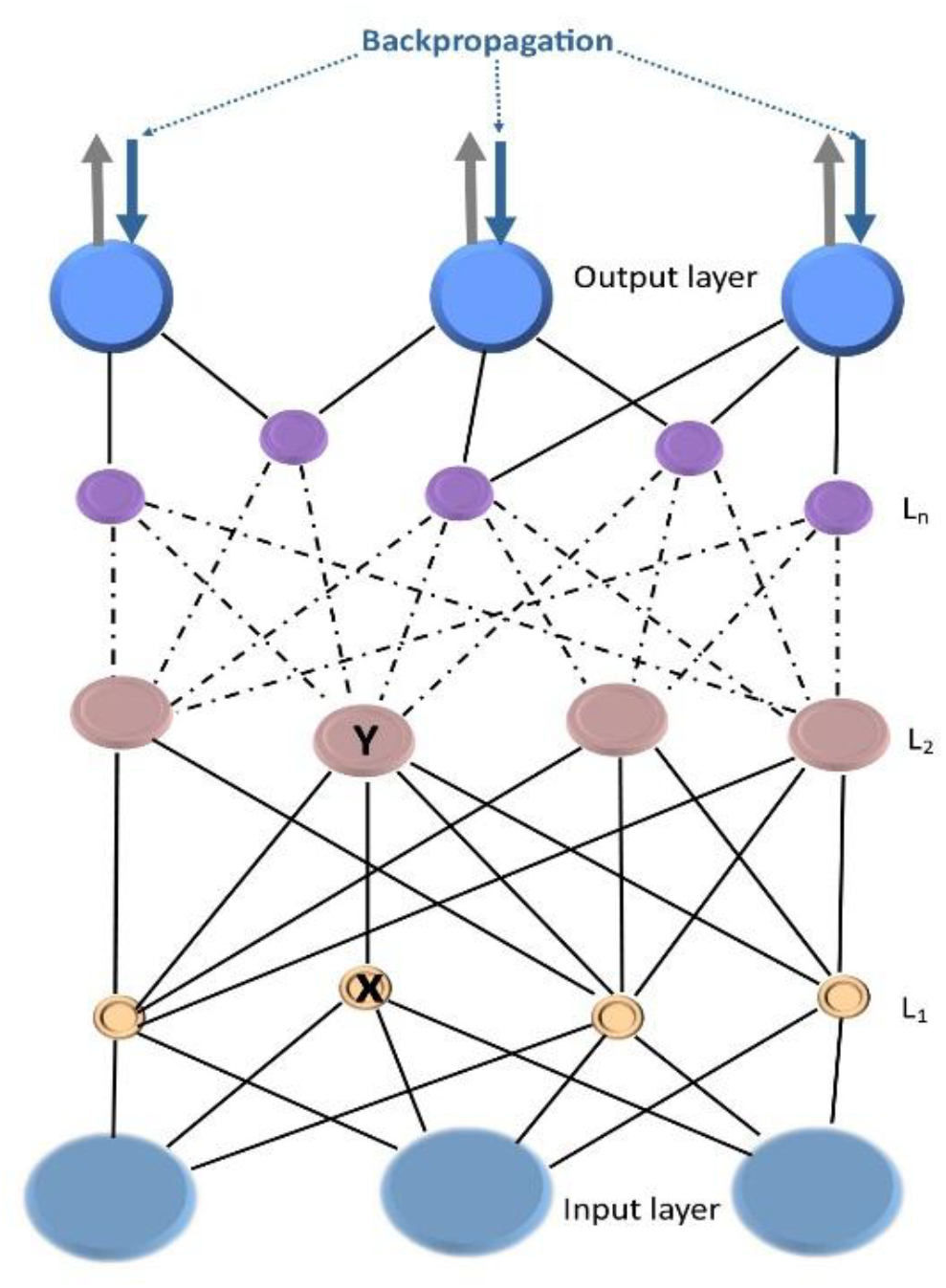
Backpropagation.

The iteration of the chain rule for derivatives is formalized by consecutively applying the following formula. Extending to *n* layers, the generalization to *n*-functions, if *y* = *f*_1_{*f*_2_(… *f*_*n*_(*x*), … )}, the derivative is the product of derivatives of each function layer, L_1_, L_2_, …, L_n_. The series of hidden network layers, L_1_, L_2_, …, L_n_ are minimizing the error and refining the identification of the object or natural language processing. The *n*-functions are evaluated hopping from one layer to the next at the inner function’s value

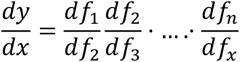

However, a weakness of the traditional chain rule is that it does not easily iterate. To find higher order derivatives (*n-*th derivatives) of composite functions, one must use *Faà di Bruno*’s formula which generalizes the chain rule to higher orders. *Faà di Bruno*’s combinatorial formula can be stated as

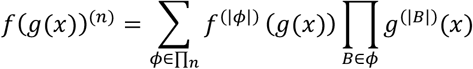

where *ϕ* ranges over the set ∏_*n*_ of all partitions of (1, …, n) recurring ^35^. For each such partition *ϕ*, B ranges over the blocks in

*ϕ*. If the derivative is the product of derivatives of each function layer, evaluated at the inner function’s value.

#### 4. Second stage, reviewing and minimizing errors by backpropagation

The backpropagation reads from top down on Fig. 1 to minimize errors between predicted and actual outputs. When trained on documents, autoencoders produce codes that enable rapid retrieval^5^. For fine-tuning, the error function^5,20^ used was the “multiclass cross-entropy error function”

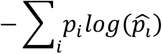

To perform those near-indefinite backpropagation or feed-forward propagation looping, AI computers require immense, rapidly increasing computing power. Data center energy demand is projected to reach 68 GW by 2027. In contrast, writing source code to formalize mathematical equations representing the human body or brain physiological processes require complex and long algorithms (in C++ or else). Instead, ML and Deep learning requires only minimal source coding in Python or PyTorch while “infinite” trial-and-error process with Bayesian inference is looping in a zeptosecond (10^-21^ s). All of which require colossal amount of energy. Python and PyTorch will perform automatic backpropagation with minimal intervention. Evolution of the human brain requires the exact opposite, energy saving-minimal volume while enhancing performance. With this equation of economy in mind, the maximum evolutionary efficiency rationale for brain structure/function must optimize energy/volume/rapidity/adaptation/creativity with energy and volume as limiting parameters.

Recent CNN architectures have 10 to 20 layers of ReLUs, hundreds of millions of weights, and billions of connections between units^2^. By 2027, analysts report anticipate significant growth inflection for companies as Mobileye and NVIDIA will be using such CNN-based methods in their upcoming vision systems for EV cars such as Tesla. Most major technology companies, including Google, Facebook, Microsoft, IBM, Yahoo!, *X* and Adobe initiate R&D projects in the performance of CNN-based vision systems ^2^.

#### 5. Machine learning–like mechanisms mirroring human brain processing

Reasoning in style with ML, rather in a less structured manner may provide hints on how the brain may work. The trial-and-error process, with Bayesian inference, seems to be ubiquity learning mode in child development^30^ and adult^36,37^. Many studies, both electrophysiological and lesion based, indicate that some visual areas, such as those in the temporal and parietal lobes, are involved in a higher level of information processing than that mediated by occipital areas such as VI and V2^38^. Hierarchical organization in the primate visual system is suggested by existence of connectivity patterns, with forward or feedback pathways^39^. The possibility that the visual cortex might operate by a strictly serial processing scheme can be ruled out just from knowing the multiplicity of connections per area and the near ubiquity of reciprocal connections between visual areas and temporal or Broca^38^. However, flexibility and non-hierarchical features were also observed in the brain. Feedback complexity with modulatory signals the same as convolutional and ReLU are also present. Parallel processing and shortcuts, direct “skip-layers” are bypassing dedicated intermediate associative cortex areas, indicating a more networked architecture. As corrections are allowed in DL, task-dependent attentional demands and behavioral context may reorganize the functional hierarchy. Numerous studies comparing performance of human brain to ML, based on different techniques were implemented. ML modalities are in constant progress. Direct single-cell electrodes recorded 395 neurons connected with prefrontal, inferior temporal cortex, playing a role with visual interpretation and behavior^40,41^. Another team compared ML techniques to visual cortex (V1, V2, V4) and IT^42^. One of the most striking aspects of visual responses in other areas, both the prefrontal area, PF, and inferior temporal, IT, is how quickly they occur. In the IT, neurons begin responding approximately 100 ms after a stimulus onset, while response onset in the PF is typically only slightly delayed^41^. Although 100 ms might seem relatively long, it is brief when considering the many stages of processing involved (see Fig.4). Visual information travels from the retina to the primary visual cortex (V1) through the thalamus, then undergoes additional processing in areas V2 and V4 before reaching different regions of the IT and ultimately the PF. In the next sections, we will learn more about other hints on how ML may be deeply embedded in the brain. In ML, stochastic gradient descents, SGD, are usually computed using backpropagation^1,2.^

The past decades, systematic well-designed studies have thoroughly investigated how brain worked based on ML and neurophysiological models, attempting to mirror the brain and identify parallels^3,43-47.^ Some authors modelled neocortical pyramidal neurons and suggested that “a critical, missing component in the current models of the neurobiology of learning and memory is an explanation of how the brain solves the credit assignment problem”^43^. Hereby, why well-designed studies end with such striking negative conclusion? The conclusions of those studies were likely right. Artificial intelligence was initially created based on the brain as a model for DL, backpropagation and other techniques^1,2^,5. Trial-and-error and Bayesian inference seem to be a universal mechanism of learning in children and in adults^30^, and so well replicated by DL and backpropagation^1,2^. Therefore, how does such a cleavage exist between AI and neuroscience when trying to match the underlying mechanisms? Geoffrey Hinton writes: “This makes the knowledge contained in the program or the weights immortal: The knowledge does not die when the hardware dies”^3^. Then Geoffrey Hinton continues: “These parameter values are only useful for that specific hardware instance, so the computation they perform is mortal: it dies with the hardware. The separation of software from hardware is one of the foundations of computer science and it has many benefits. It makes it possible to study the properties of programs without worrying about electrical engineering”^3^. All those deep analyses, statements and conclusions paralleling AI to the brain are genuinely true. Indeed, there is no mortal computation, the trial-and-error process, stands by indefinitely valid, both in terms of AI computing^1^ and human learning^30^.

Hereby, where is the gap, the discontinuity between both, the missing link? How could AI and the brain work be achieving success in their own way? Human biology and physiology are a “wall” of high dimension data, peppering smoke screens with added (essential) complexity and masking the laws of physics in the background. Mathematical description is essential to sort the physiological and hidden physical multilayers, and unveil potential corresponding CNN, backpropagation, supervised learning, reinforcement learning, self-supervised learning, feed-forward propagation, generative prediction, autoregressive prediction, etc. it does not mean that because processes as not obvious, they are not present in the background. Missing links to neuroscience are numerous, electricity propagation in conductor wiring and circuitry (transistors, chips, capacitors, etc.) is absent in the brain. Evolution of the brain focusing on minimizing space, energy and weight has developed marvels of adaptation and ingenuity. The goals of the human brain in the environment are not the prediction of a country’s GDP and interest rates day by day. The vectors of information and propagation of the action potential in brain neurons (next Section) are dramatically different than in the most powerful AI data centers. Some mathematical heuristic models described here were tested in humans^16,22,24-26,48^ some were not, albeit nonetheless essential. The equations modelling underlying ventricular ejection fraction, exchanges of gases, ventilation and brain protection in outer space are unseen and cannot be directly read in the human body^16,24-26,49^. However, the hidden laws of physics governing physiology and biology are present. Biology and physiology are adding dimensions to the multilayers of neural networks providing further economy of volume, energy and weight. The Hadamard gates models fundamental single-*qubit* logic gates and analyzes the transition of superposition of states, transient states between different vectors of information, known ones (ion channels, *Na*^+^, *K*^+^, *Ca*^*2+*^) and others yet unknown. This is essential to detail this path although it has not yet been tested in the human brain. We designate those unknown elemental vectors of information or basic units of quantum information as *qubit*. Although, at this stage of the demonstration we are only hypothesizing that they could be assigned binary values as 1, or 0.

### A-II. Charge-trap flash (CTF) technology as a model of information storage

#### 1. Quantum tunnelling, electromagnetic fields, and microprocessor memory cells

At this stage of our investigation, it is essential to draw upon established functional analogues. The functional width of the memory cell must be in order of less than 80 nanometers; deviations beyond this range render the device inoperable. At this scale, quantum mechanical effects become dominant. Modern flash memory consists of massive arrays of such cells, stacked in hundreds of layers and organized into tens of thousands of rows and columns. These nanoscopic units, known as charge trap flash memory cells, store information by trapping electrons. Within each cell, varying quantities of trapped electrons represent three bits of data, which can be retained for extended periods. Inadequate electron confinement would result in data corruption, leading to the loss of stored images, videos, and files.

To inhibit electron leakage, the charge storage region is enclosed by dielectric materials that act as insulating barriers as illustrated on figure 2 (adapted from^50^). These dielectrics are non-conductive and prevent electrons from freely traversing the barrier under classical conditions. From a classical perspective, an electron behaves as a localized point charge with a discrete energy level and cannot overcome the dielectric barrier without substantial energy input, which would damage the structure. However, quantum mechanics describes the electron not as a point particle, but as a probability density distribution function that defines the likelihood of its presence in space^51,52.^

**Fig. 2.**
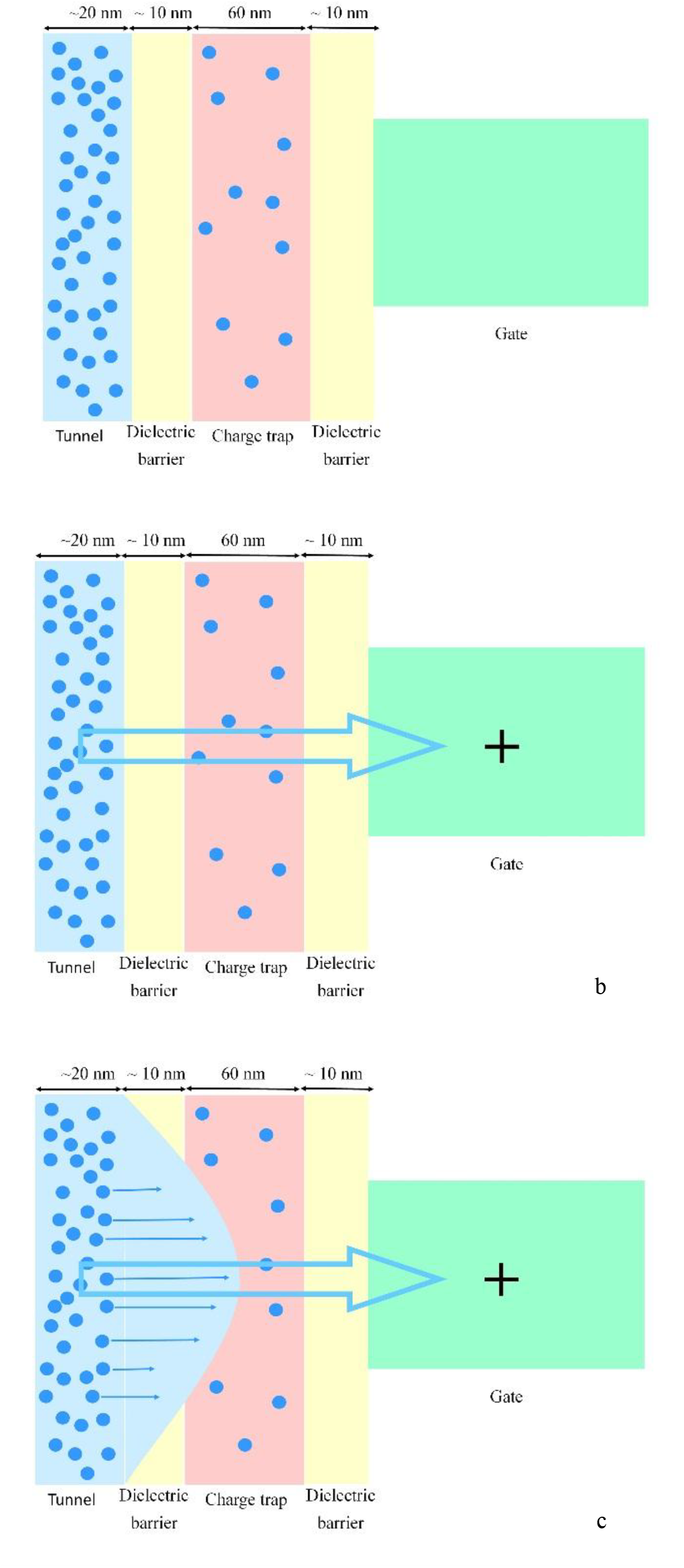
Quantum tunneling. a). Charge trap flash, CTF. b) and c). In the nanoscale (nanometer), electrons move from the tunnel when an electromagnetic field is applied and are trapped in the CTF. This is how data is stored in modern 2026 3D NAND SSD, solid-state drive to stack large number of non-volatile flash memory cells (NAND).

**Fig. 3.**
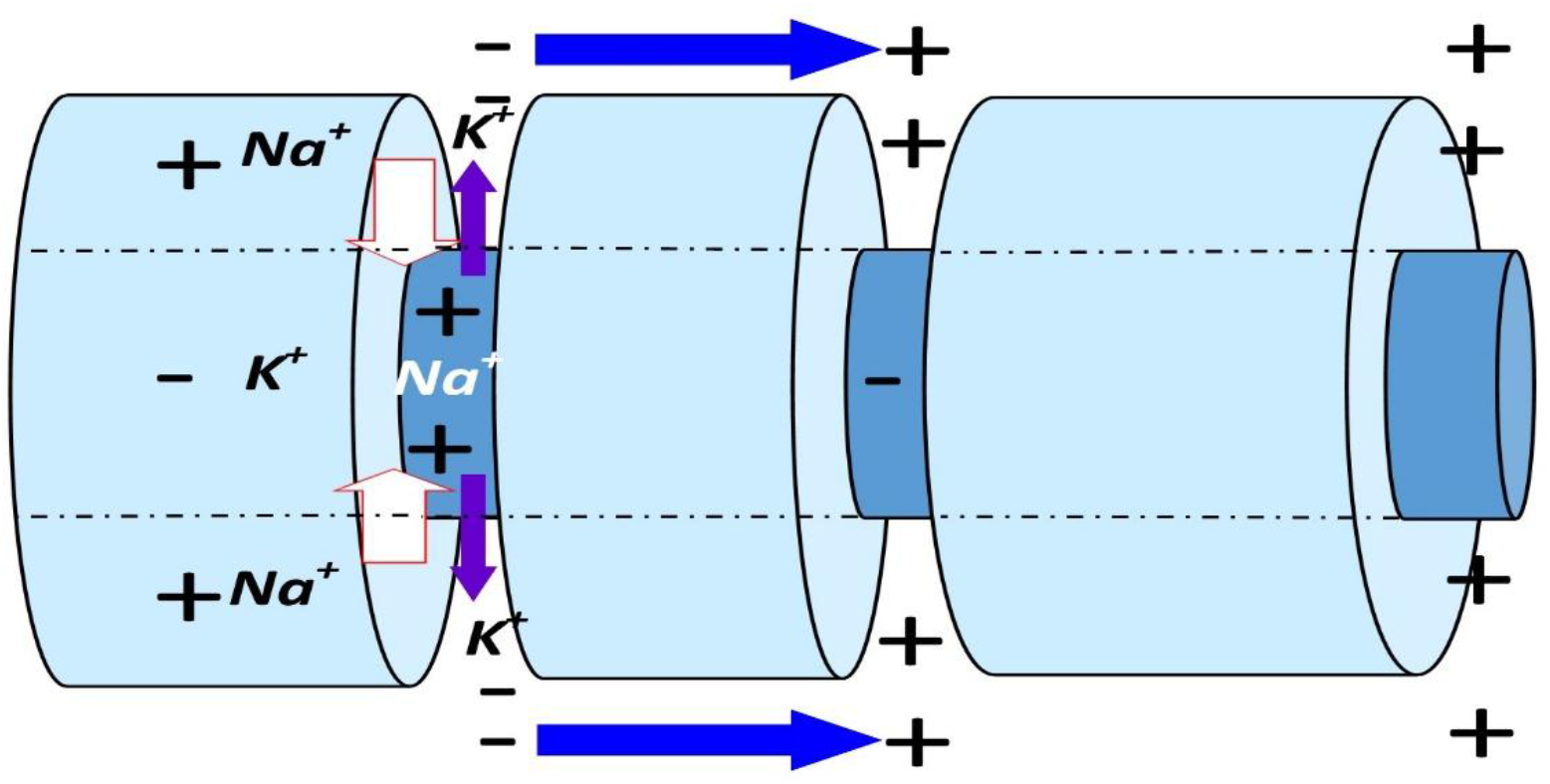
*Transversal transfer* across cellular membrane: ion channels *K*^+^/*Na*^+^ pumping (ATP-energy-system) driving ions against their electrochemical gradients. Excess of ions and net charges in a compartment are denoted by element-symbols (*K*^+^, *Na*^+^) and polarities (-, +). At rest: -70 mV relatively-negatively charged intraneuronal/axonal compartment. At depolarization, AP: +30-40 mV intraneuronal, whereas extraneuronal space becomes negatively polarized. *Longitudinal jump* of electromagnetic potential along the axon; negatively charged particles are transferred to the next node, aka *quantum teleportation* in quantum physics (marine-blue arrow). Myelin wraps: light blue cylinder; Axon: internal dark blue cylinder. Depolarization: *Na*^+^ is transferred intraneuronal (white arrow); *K*^+^ goes extracellular (purple arrow). The *NET* exchange results in negatively charging the extraneuronal space because more *Na+* are transported inside the axon. This phenomenon is important for understanding subsequent mechanisms underlying the engram formation.

When a positive voltage is applied to the gate electrode, the resulting electric field attracts the negatively charged electron probability distribution cloud from the channel toward the charge trap region^53^. If the dielectric barrier is sufficiently thin and the applied electric field is strong enough, the electron’s probability density extends across the barrier, creating a non-negligible likelihood that the electron will appear on the opposite side within the charge trap^53-56^. This process, known as quantum tunneling^54-56^, allows electrons to pass through a potential barrier rather than surmount it. Each time data are recorded, this tunneling mechanism is utilized to write information into the non-volatile charge trap memory cells. The precise relationship between dielectric thickness and gate voltage is determined using quantum mechanical models^54-57^.

One of the most remarkable aspects of this technology is the extreme thinness of the dielectric barrier. These structures represent some of the smallest features ever produced through large-scale manufacturing. At only 50 – 80 nm thick, the dielectric cannot be made substantially thicker without necessitating higher operating voltages, which would increase the risk of device damage^58,59^. Conversely, reducing the thickness would significantly raise the probability of charge leakage^58,60^. Despite these constraints, billions of nearly identical charge trap flash memory cells are reliably fabricated and integrated into devices worldwide. Over time, however, these charge traps gradually lose stored charge, with data retention typically lasting on the order of a decade under inactive conditions^61^. Additionally, the memory cells are subject to a finite number of write and erase cycles, detrapping increasing numbers of electrons, which ultimately limits device longevity^61,62.^

#### 2. *Transversal transfer* of charged ions across the axon, ion channels pumping *K+* and *Na+*

The architecture of the brain and AI-computers are both energy cost-driven and energy-savvy by design. The human brain achieves vastly superior energy efficiency, performing around 10^18^ mathematical operations per second on just 15–20 watts of power, whereas computers consume about 500 watts to carry out only about 10^12^ operations per second. Transfer of information and energy are intertwined. In the brain, most energy is used to reverse ion influxes generating excitatory postsynaptic currents (EPSCs) and action potentials^63^. EPSCs was restricted to minimize energy consumption, although kept at sufficient level not to jeopardize information transmission^63^. In biology, it is inferred that there is “one bit in one neuron spike”^63^, of the action potential, AP, which may be considered as an extended mesoscopic approximation of several yet undetermined elemental or quantum information unit, or *qubit*_*Dep*_, propagating depolarization. Despite being “mathematically tractable,” the one-neuron-spike-model likely relies on simplifying assumptions about synaptic states and transition dynamics. In the case of oligodendrocytes myelin sheath insulation, and Ranvier nodes, *qubit*_*Dep*_, may be virtual and initially created by voltage-gated ion channels and ion pumping. An electric current, *per se*, consists of electrons moving from one place to another in a conductor, usually in response to an electric field, which does not seem to be the case of the neuronal oligodendria-myelin-sheath insulation. Association of myelin sheath-Ranvier nodes provides remarkable insulation and rapid signal transmission in independent micro-circuitries packed in limited intra-cerebral space, hereby excluding shortcut commission errors. Energetic cell-induced node pumping generates an electric field from node to node. The original, elegant and concise mathematical heuristic model of transmembrane ionic transport at the node with the experiment backing the results was described by Hodgkin and Huxley^64^. Thereafter, much work was related to the *transverse*-ionic transport across membrane at the Ranvier Node^65^.

#### 3. *Longitudinal jump* of electromagnetic potential along the axon, quantum-charged particles transferred from node to node

The *longitudinal*-*jump* from node to node remains elusive and conceals direct identification of the elemental vector of information in such micro-electromagnetic field. In quantum physics, this is also known as “*quantum teleportation*”. The physical particle does not move but rather transfers its exact quantum state (information) to another distant particle using entanglement. In physics, different supports exist for *qubit* generation, such as neural spin networks^66,67,^ *J*-coupling^66,67^, electron spin^66,67,^ spin-orbital coupling^66,67,^intra- and inter-molecular spin networks^66,67,^ electron number (charge), photon, vibrational *qubit*, or others. Yet none of these theories has gained traction, although, in the universe of physics we are immersed in, there must be one elemental starter, yet to be discovered. Based on the Heisenberg Uncertainty Principle, the particle’s exact position and momentum (spin) cannot be measured simultaneously rendering intractable this type of measures, especially in complex biological spaces. In this case, the argument of quantum physics that absence of evidence does not mean absence of objective physical reality, may also apply to neurophysiology. Experience of protecting the brain in the physical space environment with absence of gravity and pressure led scientists to re-thinking the entire biology^24-26^. Precision and sometimes instantaneous timing output relating to specific input (input ↔ output) may also be a hint of underlying elemental, yet unidentified vectors of information, quantum information unit, *“qubit”*, in the brain. Such fast reaction times require an immediate output transmission. AI models such as multi-synaptic firing (MSF) neuron models, leaky integrate-and-fire (LIF) neuron modelling networks of biological neuron models, spiking neural networks (SNNs) or rectified linear unit (ReLU)^2,31^ function attempt to mimic neuronal AP dynamics^68,69^. It is assumed in those AI programs that electric charges flow through a conductor, the circuitry of the system. The elemental vectors of information, the electric signal propagates near the speed of light because the electric field propagates instantly, causing all electrons to shake in a simultaneous and general drift. However, individual electrons, bumping through atoms, move very slowly at a “drift velocity” of only a few ∼mm.s^-1^. While supporting rapid physiological response times and output in biological neuron AI models, time-consuming processing in associative cortical areas could not be recreated by the computer. In AI computing, a classic electric current “flows” electrons, along conductive wires, components, capacitors, transistors and microprocessors. Quantum AI computers’ R&D involves neural network circuitries using specific atoms such as rubidium (Rb), cesium (Cs) and erbium (Er) atoms. All of which likely won’t apply as a direct model for brain’s elemental information vector. Besides, AI developers, facing a wall of high-dimension data, designed a smart, elegant and efficient approach of modelling information processing based on “deep learning” and structuring the data in a cascade of classifiers staged in a multi-layer neural network^2,4,5,20,21,70.^

### A-III. Subdimensions of the brain

Categorization of sub-levels is necessary to analyze two distinct domains, biological macroscopic measurements and quantum-scale mechanisms.

#### 1. Organic Brain *vs*. Inorganic AI

Advanced microprocessors can utilize as many as dozens of billions of transistors. Despite being instrumental, the focused approach detailing cells by cell types, and lower dimensions, lead to a level of complications which may be different from one year to the next in one individual and prodigious across individuals. Despite being instrumental, the focused approach detailing cells by cell types, and lower dimensions, lead to a level of complications which may be different from one year to the next in one individual and prodigious across individuals.

The human brain contains approximately 87±8 billion neurons, with other cells (85±10 billion astrocytes, etc.)^71^ and 100 to 500 trillion synapses. This approximately leads to the total number of random combinations solely for the connections and long-distance white matter association tracts and synaptic connections (see dimensions MAN & MIN-1, Fig. 4a) of 5 × 10^14^ × 87 × 10^9^ = 435 × 10^21^. In the executive approach, additional sub-level dimensions (MIN 2-5) must yet be counted with supplementary 10^n^ possibilities at each dimension, in other words: 435 × 10^21+n2+n3+n4+n5^. This demo highlights the necessity of a dual approach, focused and executive, further preventing the odds of biological infinitely small high-dimension. A risky aspect of the executive approach is to remain too close to (quantum) microchips and far away from the brain biological setting. Attempts are being made to match both settings^37,72-74^. Another view of the question is that the multi-dimensional network architecture is multiplying itself by a minimum of 435 × 10^21+n2+n3+n4+n5^ similar systems. A definite structure of the system of the Matryoshka dimensions is necessary.

**Figure 4.**
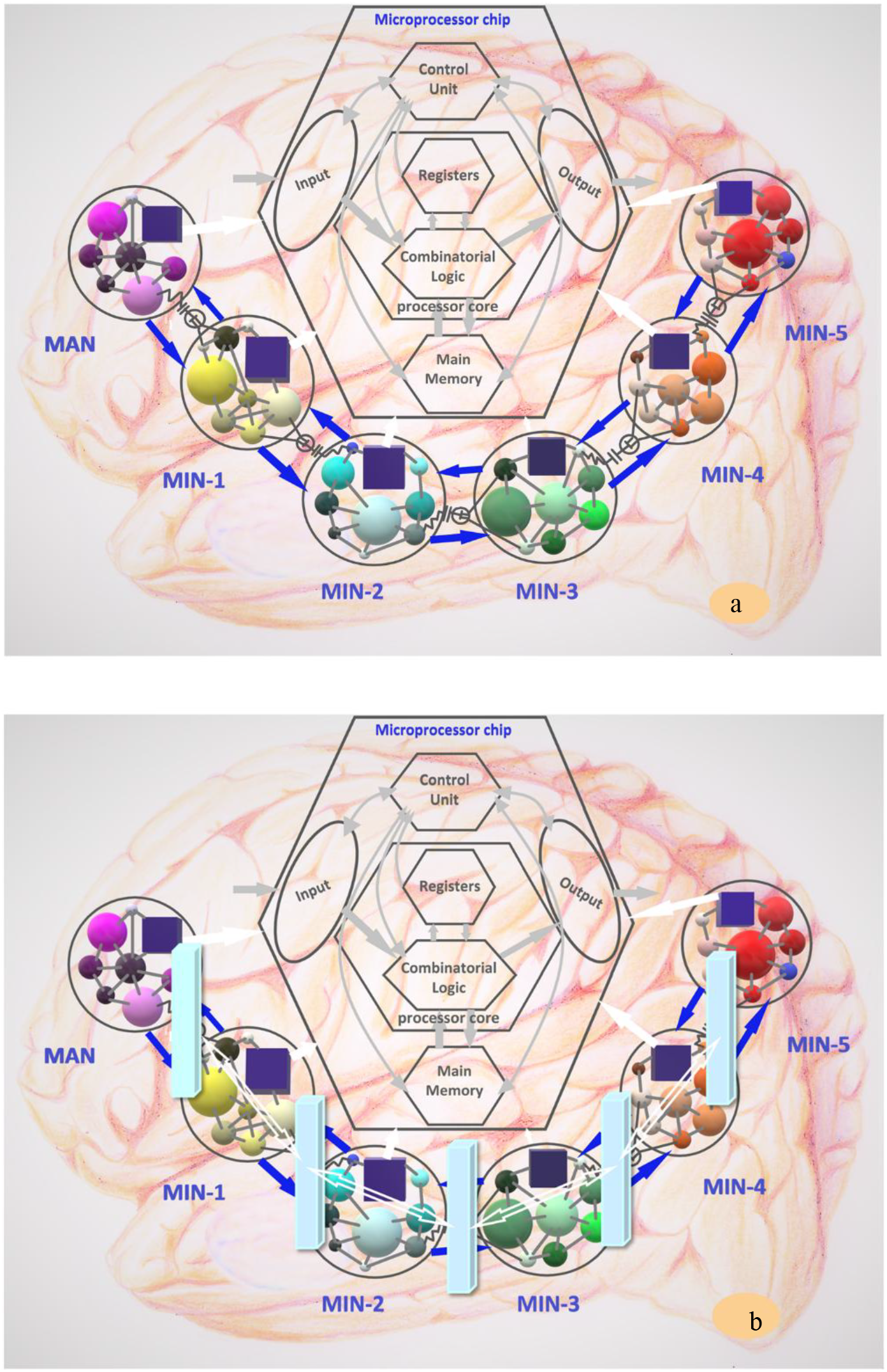
a). Multi-dimensional network. Processing of information at each dimension. ANSI electronic symbols. We intentionally sketch a microchip-like structure at each dimension. See text for details. b). Functional, physiological and anatomical barriers (light blue bars), do not strictly match the gradient of multilayers neural networks.

Yet, in the case of the brain, just looking at Fig. 4a (MAN) dimension seen on diffusion MRI^75,76^ as *3D* visualizations get more impressive, reflects the colossal complexity of the Matryoshka dolls brain network system. Across all brain dimensions, namely MAN, MIN-1, MIN-2, MIN-3, MIN-4, and MIN-5, the number of combinations amounts to zillions of options (Fig. 4a). The organic world of the living realm possesses inherent built-in obvious advantages, such as plasticity, regeneration, flexibility, and complexity, over the inert inorganic world. While visualization to assess all fractal heuristic mechanisms across two dimensions can be modeled, they are not directly accessible in real-time due to insufficient resolution. Therefore, the focus lies on computationally evaluating and describing quantitative physiological outcomes, time, and speed. The simplest models utilize basic neural mass or mean-field models to capture changes in mean firing rate, while the most advanced models employ a dynamic mean-field model derived from a proper reduction of a detailed spiking neuron^77^.

#### 2. Modelling elemental “nanoscale” units of information across sub-dimensions

Despite being instrumental and essential, the focused approach detailing cells by cell types, and lower dimensions, lead to a level of complications which may be different from one year to the next in one individual and prodigiously variable across individuals. This is exposed to the odds of high dimensions. In the brain, the information must cross structural and functional barriers where the *information-unitary or one qubit* may be using a different vector in each dimension. In AI computers, ML relies on regular electric current with electrons as information vectors across multilayers of artificial neural networks. In contrast, information along the axon travels much slower than electricity in a conductor or a circuitry. Modelling heuristically the passage of information vectors across biological structures of the brain will be essential to unveil some potential hints and mechanisms of the brain described in the next sections. However, those vectors of information are yet unknown.

The mathematical modeling approach is defined by a “quantum feature map”^51,78,79^ on *n-qubits* generated by the unitary^79^. The data is mapped non-linearly to a high dimensional space, the quantum feature space^79^. For a single *qubit, U*_*ϕ*_(_*x*_) = *Z*_*x*_, is a phase-gate with angle x; where 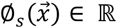. We postulate that there may be two types of *qubit*: 1). *qubit*_*Dep*_ (depolarization) vectors of the action potential along the axon; and 2). *qubit*_*Sub*_ elemental information vectors within the cell. We may envision the transfer of *qubit*_*Dep*_-information into the *qubit*_*Sub*_-information at the intra-cellular sub-levels following the classic mathematical modeling approach as defined by a feature map^51,78,79^ on *n-qubits* generated by the unitary^79^:

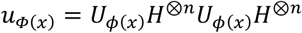

where H represents the conventional Hadamard gate^79^, and where

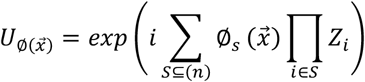

*Eq*. (6) is a diagonal gate in the Pauli Z-basis^79^. The coefficients 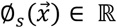 are used to encode the data × (x ∈ Ω subset of ℝ, ℝ is the set of real numbers)^79^. This theory is valuable to analyze and quantify the transformation of *qubit*_*Dep*_, into various types of *qubit*_*Subi*_ (i = 1, …, n), supporting the conversion *qubit*_*Dep*_-information to the *qubit*_*Subi*_-information in sub levels of the micro-network (Fig. 4a). From the initial “on” or “true” state, *qubit*_*Dep*_, represented by |1⟩, there is a superposition of the standard “off” or transient “false” state, *qubit*_*Subi*_ (ion channels, *Na*^+^, *K*^+^, *Ca*^*2+*^ or other intracellular-nuclear vectors), represented by |0⟩, at the transition state to another “on” or “true” state, *qubit*_*Dep*_, represented by |1⟩. In some other words, the initial “on” state, *qubit*_*Dep*_, represented by |1⟩, operated as described in Fig. 5, goes through conversion of information to a sub-level of the micro-network, from MIN-1 to MIN-5 (Fig. 4a) via transient “off” states, *qubit*_*Subi*,_ |0⟩, prior to returning to the initial “on” state |1⟩, same neuron (feedback loop) or a different neuron, with vector *qubit*_*Dep*_ (Fig. 5). The initial and final “true” states |1⟩ may be viewed respectively as “sensory input” and “behavioral output”. The transient “off” states, *qubit*_*Subi*,_ |0⟩, may be considered as avid energy-dependent processing of information in the visual areas, V_j_ (j = 1, 2, 3, 4, …, n), of the occipital cortex and visualized by BOLD fMRI voxels. Referring to ion channels, *Na*^+^, *K*^+^, *Ca*^*2+*^ participating in the exchanges is crucial. The presence and the role of ions channels is instrumental to eliminate several other implausible biological models. It may suggest hints on how ML and deep learning may fit the brain and on memory storage for long-term potentiation (LTP). This will be discussed in next sections.

**Fig. 5.**
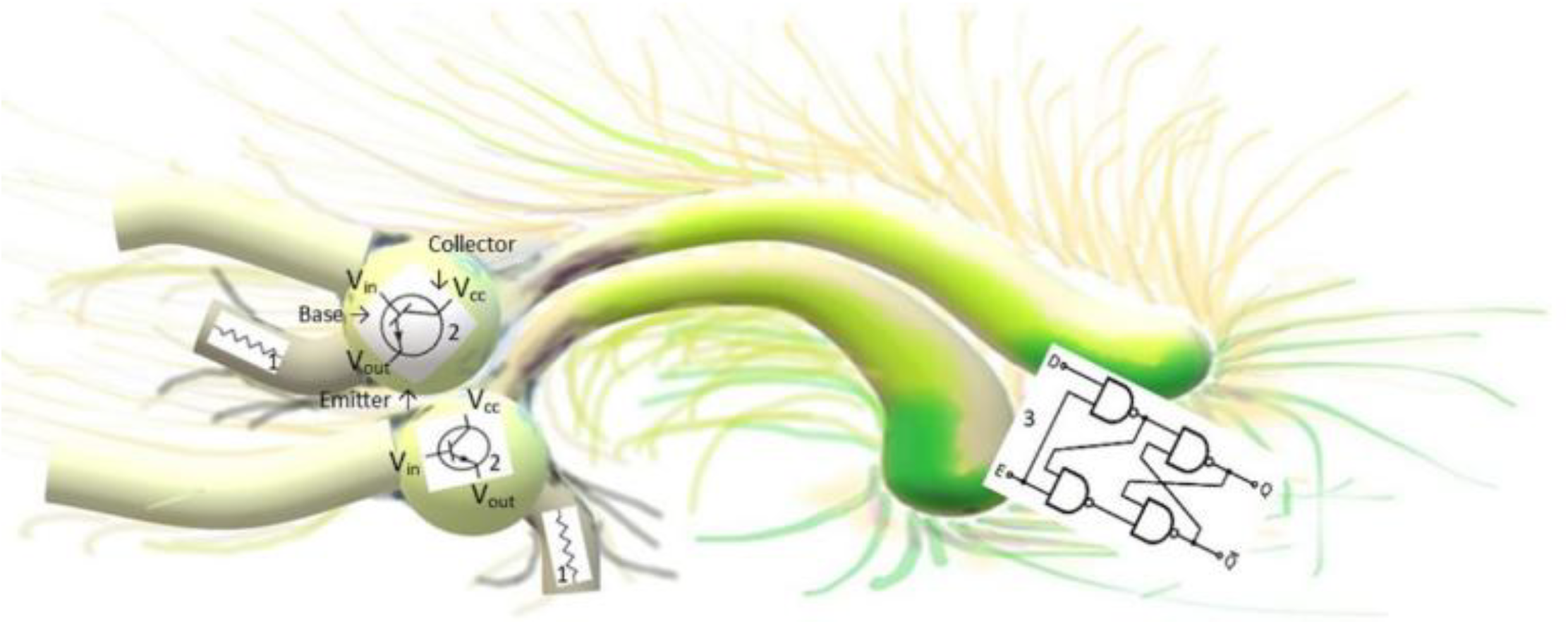
Memory Storage in an array of flip flops & logic gates in a microprocessor-like. This is a potentially most simple case in dimensions MAN & MIN-1. Resistors, transistors, and microprocessor are respectively illustrated by 1’s, 2’s, and 3.

#### 3. Multi-layers and multidimensional brain modelling

In contrast to AI computers, it is brilliant that the human brain’s action potential, AP, can be triggered anywhere and autogenerated by its own energetic system, without a generator-battery. This provides flexibility and energy saving. In most neuronal types, Aps are initiated within the axon initial segment, or the first node of Ranvier in myelinated axons, which is usually associated with a hotspot of *Na*^+^ channels^80^. From the AP-initiation site, the nodal *transverse*-ionic transport across membrane expands by *longitudinal-*propagation from node to node along axonal axis during spiking^80^. Two distinct physical-chemical mechanisms, *transversal* and *longitudin*al, are underlying the AP. The AP-initiation site expands along the axon during spiking depending on several variables such as axonal resistance, system capacitance, density of voltage-gated channels, and internodal intervals^81^. All of which regulate the conduction velocity and timing of APs, measured by patch clamp. The AP seems to be accompanied by mechanical axonal waves creating fast and temporary changes on radius, pressure, optical properties, release-subsequent absorption of small amounts of heat, and axon shortening^82^. In another sub-level of high-dimensional spaces, single-neural cell transcriptomics, progress was recently made to unveil the link between transcriptomic and morpho-electric descriptions of cortical neurons^83^. Neuronal firing patterns differing between cell types could be associated to specific ion channel gene expressions, including voltage-gated channel densities^83^. The analogy might be a tree with ramifications of cortical cell types described tending to continuous manifolds^83^ which may be stretching like an Euclidean space ℝ^*n*^. The temporal location for memory processing in a section, although to date, no elemental micro- or nanostructure has yet to be identified as post-processing potential mass storage. Grid cells, in the entorhinal cortex, are a grid-GPS-like coordinate or navigation system, with specialized neurons firing in hexagonal pattern, enabling the human brain to map the spatial environment and objects^84^. Perception of the surrounding environment or virtual world of a computer screen elicits spatial memory in order to localize the characteristic configuration of the scene on display. Perception, motion or navigation in a three-dimensional space are closely related to spatial memory requiring permanent visual tracking of the direction and distance from reference points, or landmarks, and their integration by hippocampal and entorhinal network mechanisms underlying grid cells executive mapping^85^. The grid cells are thought to encode the conceptual space, and not just the physical space^86-88^. Place cells (hippocampus), are participating in spatial navigation and memory about important or unique places; possibly receiving input from grid cells^89-91^. This temporal navigation system is densely connected with visual areas V_j_ (j = 1, 2, 3, 4, …, n)^40,41,92^. The sensory integration enables efferent responses from visual input to projections to posterior-parietal Brodman areas for motion or to Broca and Wernicke areas for speech. Our view of this integration and communication must be flexible and not strictly related to anatomical, functional barriers, nor closely associated to well-defined structural brain-multilayer neural network.

#### 4. Multidimensional approach to disentangle neurobiological high dimension spaces

Gradual reinforcement encodes cognition across multi-dimensional brain networks from *macroscale* long-distance white matter association tracts^85^ to *microscale* long-range promoter-enhancer interactions governing interactions which control gene expressions, such as *GRIN2B*^*93*^. All these different dimensions can be illustrated as Matryoshka dolls, with each dimension promoting a very specific function at its own fractal scale. The mechanism involves multi-way communication, with feedback sent both by the sender and receiver, upstream *and* downstream. This multi-level scaling and communication significantly impact cognition and working memory. This multi-dimensional system combines protein transport, molecular pathways, histone acetylation, gene expression and transcription. To be a working system, all sub-level micro-dimensions or neural network layers should be inclusive, even at micro-dimensional level, yet unknown or not validated by known physical and biological support. Invisible in our own perception does not mean non-existing^94^ and a working model of the brain must be inclusive. In AI computing, dimensionality reduction via neural networks simplifies the classification, visualization, communication, and storage of high-dimensional data^5^. Such dimensionality reduction is certainly possible in the brain configuration. We are identifying a cascade of physiological classifiers for deep reinforcement learning modelling. Based on AI current development and physiology, we are exploring such a model in the “deep learning” section.

#### 5. Schematic of brain physiological macro-network and levels of micro-network

Based on functional roles, a multidimensional network architecture for brain function is proposed and shown in Figure 4.

0. MAN, the macro-network is supported by elemental “nanoscale” units of information, yet unidentified *qubit*_*Dep*_, and described at the macro-scale level of brain network and is defined by synaptic connections and long-distance white matter association tracts in a four-dimensional space.
1. MIN-1, first level of the micro-network is defined by the synaptic (pre- and post-) space.
2. MIN-2, second level of the micro-network is defined by the axonal system and neuronal cytoplasm.
3. MIN-3, third level of the micro-network is defined by the nucleus (neuronal).
4. MIN-4, fourth level of the micro-network is defined by the epigenomic system (neuronal).
5. MIN-5, fifth level of the micro-network is defined by the genetic system (neuronal).

At each dimension, in terms of graph theory applied to neuroscience, we extend network topological analysis of weighted graphs to all systems from MAN to MIN-5, including small-worlds and rich-worlds (spheres & vortices)^85^.

Reinforcement of information occurs by a mass balance mechanism (blue arrows, Fig. 4a) between each dimension; can be thought as backpropagation-like physiological mechanism. For each *qubit* travelling MAN → MIN-5, there must be feedback information, in the form of at least *one qubit* (***n* ≥ *1***) progressing in the MIN-5 → MAN direction. Spike-timing-dependent-plasticity, STDP, is a form of neuroplasticity phenomenon where repetitive, precise timing pairing the presynaptic spike and postsynaptic spike determines whether a synapse will be strengthened (Long-Term Potentiation, LTP)^95,96.^ The Hebbian theory describes the synaptic connection strengthening after repeated use^44,47.^ Neuronal plasticity, synaptic strengthening, supports the apparent neuronal entropy or rather a variable architecture. Designing artificial neural networks (ANNs) does not involve writing source code for specific computations performed by the network. Instead, they draft three components: (1). Loss (error) functions; learning rules; and (3) architectures. Learning rules are guiding to update the current synaptic weights^44^. Applying multilayers or neural networks’ architecture to AI computers, with highly energy-consuming electric currents across conductors and circuitry (transistors, AI chips, capacitors tec.) mitigates the dimensions of CNN, backpropagation, feed-forward propagation etc. This is restricting the connective-architecture to electron-activation in the AI-circuitry or overlaying a blueprint of pyramidal neurons connectome (dimensions MAN and MIN-1) into the AI design. The physiology from sub-levels of micro-networks, MIN-2 through MIN-5 is exponentially adding dimensions to the brain which may be further utilized as multilayers or neural networks. All of which are at a low cost in terms of energy, volume, weight, rapidity and plasticity/adaptation/creativity potential. The brain is a marvel of ingenuity in optimizing the trade-off between parameters to be minimized and those to be maximized. Evolution led to this state-of-the-art brain. We could imagine that human evolution could have selected gathering different materials from nature and use atoms Rb, Cs, Er, atoms, readily leading the brain to an AI-computer architecture. However, readiness to instantly compute the daily GDP of a country did not seem useful to early Homo sapiens.

Attempts to reconcile the apparent disharmony between the brain and backpropagation in a ML model which amends synaptic updates^47^. Feedback connections inducing neuronal activities and locally computing differences and encoding backpropagation-like error signals in a ML model aka “neural gradient representation by activity differences” NGRAD^47^. Adding random feedback weights may provide useful teaching signals^97^. Designing an AI architecture mimicking the brain is facing the “credit assignment problem”^43,45.^ This architecture must define the multilayers of neural networks in a hierarchy of circuits to work. Adding physiological dimensions inherently present in the brain may help solving the issue of the missing link^43^. Adding the inherent physiological brain dimensions may help create a mirrored image of the brain in AI. In the next Section, we are describing 6 dimensions (i = 0, 1, 2, 3, 4, 5). This hierarchical architecture is not limiting; there may be more currently “silent” physiological layers. This physiological hierarchy may be strictly overlaying on AI architecture. Sometimes functional, physiological and anatomical barriers do not strictly match, as scientists experienced in the space program^25^. The stochastic gradient descent (SGD) across multilayers neural networks may not mirror logical physiological gradients across anatomical and functional barriers. On Fig. 4b, light blue bars sketch some of the existing physiological; this is a simplified model, albeit existing additional functional barriers may also be relevant to ANN and DL.

Emancipating from the Moravec’s paradox^8-11^ and reaching out to artificial general intelligence, AGI, or “world models”, a hypothetical form of AI designed to mirror the overall human cognitive abilities including imagination, creativity, and consciousness is one of the current ambitions of the field. Attempts were made to human-like systematic generalization of AI^98^. This was the subject of a discussion in January 2026, at the world economic forum (Davos, CH), between Demis Hassabis and Dario Amodei. Discussions were also about advancing plans for space-based AI data centers cooling down in the extreme cold outer space using starlink V3 satellites grid (SpaceX and Starcloud with NVIDIA GPU-enabled satellite).

For each dimension scale, a microprocessor chip is represented by a purple rectangular box and its function is explained at the top of the diagram. To simplify the illustration of functional features of signal treatment and processing, we use ANSI electronic symbols. A caveat to this formal electronic description is that elemental quantum information units are yet undefined. This systematized description is extensively used in neutral-atom quantum computers (Rb, Cs, Er, atoms), where their electronic states and electron spin are leveraged to create *qubits* and perform quantum operations, which are fundamental to developing advanced AI systems. Although supported by different vectors, this physical architecture may also fit the brain, describing the lag time, various gateways^99^, molecular pathways^100-103^, transcription factors^93^, gene up or down-regulation^93^. Hereby, Bayesian options^37,72,104^ are also necessary for the storage of information resulting from signal treatment/processing^105^.

Correlating brain network activities in hubs and white-matter tracts is based on blood oxygenation level–dependent (BOLD) functional MRI (fMRI), which is assumed to mirror or reflect in part or whole, the neuronal activity^99^. BOLD fMRI is designed to measure changes in the heterogeneity of the magnetic field within each small volume of tissue, resulting from local changes in metabolism and blood oxygenation. These changes in local metabolism and energy consumption modify the ratio of deoxy- and oxyhemoglobin which have different magnetic properties and can be detected by BOLD fMRI^106^. Notably, BOLD signals had significantly higher variability than neuronal activity, indicating that human fMRI underestimates the reliability of its relation to neuronal activity^107^. These limitations of fMRI are not related to physics and are unlikely to be resolved by further sophistication and power of the scanners; instead, they are due to the circuitry and functional organization of brain networks^108^. It’s important to recognize that the content of a single voxel is both neuronal and vascular.

#### 6. A very simple memory: the synaptic flip-flop-like microprocessor

This model is unlikely to constitute a general mechanism in the brain, as it would be both spatially inefficient and energetically costly. Let’s look at a demo on how a neuronal synapse may work in terms of electrical engineering. We would be assuming the vector of elemental information would be electricity propagation in conductor wiring and circuitry (transistors, chips, capacitors, etc.). All of which is indeed not the case in the real brain biological connectome micro-environment. Considering that the net result in terms of information-output may be approximately identical from engineered DL-backpropagation and neurophysiology.

As illustrated in Fig. 5, the depolarization from the action potential across the myelin sheath and the neuronal cell membrane elicit the firing rate and signal transmission, which are exclusively defined in the macro-network, MAN, and the first level of the micro-network, MIN-1. Some of the electrical firings will dissipate in energy along the way to carry on unknown functions. A lot may also be diverted and dissipates towards sub-levels of the micro-network, MIN-2, MIN-3, MIN-4, and MIN-5.

Suppose a small intensity voltage from a subliminal stimulation (Fig. 5), let’s say visual, translated into a small change in voltage, V_in_, changes the small current through the base of the transistor whose current amplification combined with the properties of the circuit means that small swings in V_in_ produce large changes in V_out_. V_cc_, is the voltage at the common collector.

Memory may be stored by electricity/firing signal with an array of flip-flops in the microprocessor (number 3) in Fig. 5. The circuit (3) has a feedback loop. Each operation created with logic gates is mapped in the microprocessor to a number called an opcode that the microprocessor recognizes as the number corresponding to that operation (1 or 0). Now, if Emitter E (enable) has voltage, Q (*e*.*g. Q* = 1) will copy D, and Q will keep its value when E is disconnected (*e. g*. 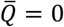). This might be a crude way to store memory at the macro-network level, MAN, and the dimension of the micro-network, MIN-1. The scale of Fig. 4’s network, yet in the dimension MAN and MIN-1, may be considered at a much smaller scale, Fig. 4a. Note that in the brain, memory may also be stored at sub-levels of the micro-networks^109,110,^and this could be done all the way down to MIN-5.

There is a duality of the brain. On one hand, the fast “electromagnetic phenomenon (AP)” one, relying on rapid *qubit*_*Dep*_-information transfer, enabling fast reaction-times (< second), life-saving capacity in evolution. The electromagnetic phenomenon (AP) is by far not as fast as electricity in AI. Although the human brain may enable fast reaction times [fast input ↔ output] through this path (Fig. 5), its processing speed cannot match AI-Computing. On the other hand, the “slow” processing brain, taking hours, days to proceeding and relying on *qubit*_*Sub*_, allowing exclusive abilities of the human brain, such as creativity, imagination, and consciousness. Reading is slow processing^111^, see next Section. This slow processing phenomenon relies on unknown elemental vectors of information. However, this does not appear to be a general model for memory storage because the volume in the brain would be excessive for a single binary information stored. AI-computers do not store in such a way.

## B. Results

### B-I. Long-term memory storage

#### 1. Inventory of existing models for memory formation

Over the past several decades, extensive research has focused on the making of LTP in the brain^112-115^. Synaptic plasticity and synaptic augmented strength have been pillars the memory making^44,47^. Enhanced synaptic strength further contributing to the induction and maintenance of LTP^116,117^. Episodic memories are thought to be encoded by experience-activated neuronal ensembles, however, the memory engrams after initial encoding remain poorly understood^118^. Furthermore, inhibitory synaptic plasticity during memory consolidation has emerged as a critical mechanism for engram selectivity^118^, Yet, synaptic strengthening alone is insufficient to account for the dynamic sharpness of enduring memories^119^. Memory consolidation involves spontaneous, brain-wide network reorganization during rest and sleep, albeit the underlying mechanisms remain unclear^120^. Evidence suggests that resting-state network hubs contribute to memory consolidation implicating a widely distributed network beyond the hippocampus^120^. Astrocyte–neuron communications further shape neural circuitry assembly and function^121^. Through rapid glutamate release, astrocytes can control excitability, plasticity and synchronous activity of synaptic networks^121^. Despite recent advances in identifying an engram, distributed in broad brain areas, the cellular and circuit-level mechanisms by which engrams are generated during memory formation remain unresolved^122^. Perception, motion or navigation in a three-dimensional space depend on spatial memory and require permanent visual tracking of the direction and distance from reference points, or GPS-like landmarks, and their integration by hippocampal and entorhinal circuits supporting grid-cell-based spatial mapping^84-86,88^,123-125. Hippocampal sequence events, referred to as “replay”, facilitate goal-directed navigation, supporting memory-based trajectory planning and guiding subsequent navigational behaviour^126^. The CA3 region plays a critical role in maintaining working memory representations in delayed-to-match sample tasks and hippocampal neuronal dynamics encode a difference between correct and incorrect trajectories^126^. Our sensory systems may collect data at ∼10^9^ bits.s^-1^ enabling remarkably fast reaction times^111^. A tennis player can be judging the speed (∼ 130 mph), the strength, future geo-localization of the ball in space, forecasting its direction, running, dodging, anticipating the body position, racket placement, optimizing the shoulder-arm lever in space, while briefly checking the angle on the other side of the net, and hitting the ball with slice effect for a winning point. All of which, accomplished in a fraction of second. In comparison, the processing speed of the brain, information throughput of a human brain was evaluated ∼10 bits.s^-1^, relatively small^111^. This was evaluated based on ∼1 bit/character in English language, the brain seems to produce an output of ∼10 characters.s^-1^ or ∼10 bits.s^-1^. Strikingly, two very different timings were necessary in two brain complex information processing. The tennis player *vs*. reader reaction times are deeply intriguing. Time boundary cells, specialized neurons in the entorhinal cortex, hippocampus, and lateral prefrontal cortex (LPFC) are internal temporal organizers of memories segmenting working-memory and navigation epochs necessary for recalling goal-directed paths^127^.

Recent studies shed light on the importance of epigenomic (MIN-5 Dimension) and genomic (MIN-4 Dimension) mechanisms in memory formation^128^. Memory encoding is associated with increased chromatin accessibility, predominantly on enhancers regions, followed by spatial reorganization of 3D-genome architecture and promoter-enhancer interactions^128^. Memory encoding induces epigenetic priming, marked by increased accessibility of enhancers without corresponding transcriptional changes^128^. Neuronal activity can also induce DNA double-strand breaks (genome-wide NGS) at specific loci in vitro, facilitating rapid transcription of early response genes, ERGs, such as *Fos, Npas4*, and *Egr1*, and repair^129^. Those neurons typically undergo energy-intensive molecular adaptations, occasionally resulting in transient genomic instability and DNA damage. The cascade of learning-induced molecular events precipitates discrete neuronal clusters undergoing double-stranded DNA damage and TLR9-mediated repair, resulting in their recruitment to memory circuits. Interneuronal perineuronal nets further stabilize memory circuits by regulating inhibitory control inputs to specialized neuronal assemblies^130^, while age-related alterations of medial-entorhinal cortex perineuronal net density and distribution, contribute to spatial navigation deficits^131^. Increased translation machinery facilitates spine remodeling, size and number, resulting in enhanced synaptic plasticity^128^.

Memory generalization and the underlying mechanisms, by which cognitive control regulates working memory storage, remain unclear^132^. Memories are encoded within neural ensembles during learning and stabilized through post-learning reactivation, yet the conditions that enable memory linking across extended temporal intervals are unknown^133^. Offline periods provide an opportunity for memories integration and retrospective across experiences. In the orbitofrontal cortex, a distinct population of neurons, the memory trace (MT) neurons, encoding action–strategy memories supports flexible decision-making^134^. These MT-neurons exhibit a higher proportion of mature dendritic spine types than neighboring non-MT neurons, a profile closely associated with successful learning.

Recent data on the dynamic and selective engram, by inhibition process shed light on memory formation and reshaping^118^. Feed-forward and recurrent excitatory synapses onto excitatory neurons exhibited short-term and long-term plasticity, whereas inhibitory synapses onto excitatory neurons displayed inhibitory plasticity^118^. Long-term excitatory synaptic plasticity combined a Hebbian term, consisting of triplet spike-timing-dependent plasticity (STDP), and non-Hebbian terms, including heterosynaptic plasticity and transmitter-induced plasticity. Mean weight strength of plastic synapses clustered according to engram cell status. Top, feed-forward excitatory synapses onto excitatory neurons^118^. Although several brain regions encode social information, the cortical circuits responsible for consolidation of memories and the nature of the information stored within the neocortex remain largely unknown^135^. Strengthening is indeed observed at synaptic connections between interregional cell-ensembles activated during learning and is crucial for memory retrieval. Despite recent advances in identifying an engram, how the engram is created during memory formation remains elusive^122^.

#### 2. Exclusion of implausible models for physiological memory storage and engram inference

In the previous section, we analyzed the significant progress that has been made in elucidating the cellular machineries underlying memory formation. This inclusive and thorough analysis of memory formation are the pillars providing deeper insight to further elaborate on physics and physiology. Indeed, the precise physiological, molecular, and physical (quantum) substrates of memory storage, and location, remain an open question^118-120,122,132^. ***It does not exist a single specific protein nor single specific gene encoding for a single specific information-memory bit***. The principle of exclusion (Michelson–Morley) is guiding to cancel implausible biological models underlying the storage for long-term potentiation of memory. Hereby, we are left with a finite scope of physiological and physical paths. Let’s cite the example the initial identification of the Higgs boson which was based on its indirect (1964)^136,137,^ rather than late most formal detection at CERN (2012). The elegant demo was the “popping up” of a “statistical excess or bump” on the curve at ∼125 GeV, only observed in invariant mass distribution. In the space program, scientists faced an unknown phenomenon, and they excluded alternative possibilities because there was no other atoms present in a free state (also invariant) in the body which could participate in the laws of physics involved in the process^16^; they could prove it with a thorough statistical analysis (maximum likelihood).

#### 3. Equipment inventory of the brain machinery for the engram

Let’s pursue the inventory of the physiological, biochemical, and physical equipment of the brain started in previous sections. A single oligodendrocyte can wrap and insulate up to 40-50 axon segments. The structural integrity of myelin layers is necessary for learning and memory^138^. Myelin layers are compacted to a thickness proportional to the axon diameter^139^. A single axon can have from 10 to 160 and up to 250 or 300 layers for high-speed transmission with an average range from 40 to 100. The thickness of a single myelin layer ranges from 7.0 to 20 nm. The dimension scale is in nanometers, approximately the exact dimensions of the charge trap, the tunnel and dielectric also equipping current AI microchips, type NVDIA, Intel, Tesla AI-5, etc., going thinner and smaller with a trade-off of damaging the dielectric membrane. On the axonal transversal slice, Figure 6a, we see the cylindrical superposition of myelin layers. The internode length typically ranges from 50 μm to 750 μm with nodes about 1-2 μm. The internode-length is sufficient to enable rapid transmission of the electromagnetic potential to the next node. The superposition of multiple, concentric, spiraled, and highly compacted cytoplasmic membrane wraps are assembled by the oligodendria (oligodendrocytes) (Fig. 6a). This spatial packed wrapping in 3D creates a disjointed multiple assembly; nodes not facing from a layer to next (Fig. 6c). This is in fact amazingly beneficial for perfect isolation emulating the memory cells of the current AI chips, collectively consisting of a massive array of cells, stacked in hundreds of layers and organized into tens of thousands of rows and columns. Individually, enabling rapid conduction with sufficient internode length.

**Fig. 6.**
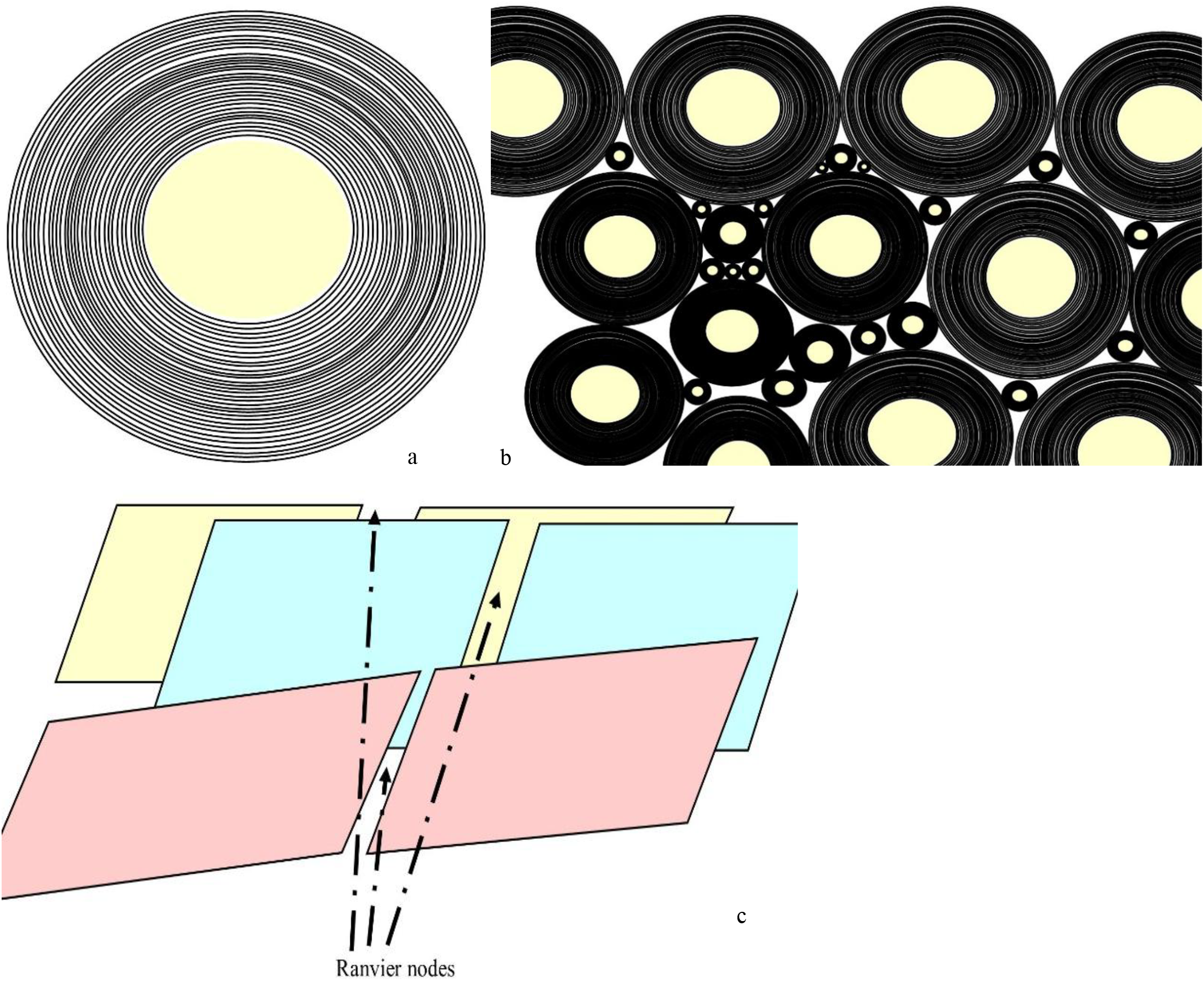
Myelin sheath. a). Transversal slice of the axon and its myelin insulation. Superposition of up to concentric 300 layers wrapping a single axon. Each one could host hundreds to thousands of niches, “memory cells”. b). Transversal slice of a long-distance white matter tract, the bundle may contain up to 200-200 million compressed myelinated axons. c). *2-D* plane of unfolded myelin layers exposing the random disjointed placement of Ranvier nodes exponentially increasing the hosting of charges ions in their memory cells. Axons within a single long white matter bundle fire asynchronously at different times.

Let’s look at the number of Ranvier nodes. The number in a single neuron varies^81^, depending on the axon length, ranging from a few to hundreds or thousands. In the human brain, for an average of 87 billion neurons (SD = 8), the total number of nodes would approximately be ranging from N = 87 × 10^9^ × 10^x^, hereby, × = (3, 4), and N = (87 × 10^12^; …, 87 × 10^13^) ≈ (90 × 10^12^; …, 90 × 10^13^) ≈ (1 × 10^14^; …, 1 × 10^15^). This would approximately be one trillion to hundreds of trillions. Figure 6b illustrates the fused layers of oligodendrocyte-insulators wrapping the axon. The myelin is constituted of phospholipid bilayer (and proteins), highly insulating. There are minute-masses of cytoplasm trapped between fused layers of cell membranes of oligodendrocytes. Those virtual spaces of relative “negative or lower pressure points” are important for adhesion and trapping (major dense line *vs*. intraperiod line).

Let’s look at the electrolytes involved in the action potential, AP. A physiological regulation of electrolytes controls intra- and extra-cellular milieus within a strict homeostatic range. The plasmatic *Na*^+^ normally ranges within 138 - 142 mmol/L, and plasmatic *K*^+^ within 3.5 – 5.0 mmol/L. In terms of electromagnetic potential difference, sodium and potassium are the two main ions involved in the axonal depolarization and repolarization of neurons (Sections *“Transversal transfer* of charged ions across the axon, *Longitudinal jump* of electromagnetic potential along the axon”). This homeostatic control is vital for the organs and cells. The loss of electrolyte balance will impact with severe life-threatening consequences to the heart, all organs including the brain, and all cells. This intra-, extra-cellular, and plasmatic electrolyte balance of *Na*^+^ and *K*^+^ applies beyond the blood brain barrier, encompassing neurons, oligodendria, glia, and microglia. This is critical to maintain the normal negative resting value of neuronal membrane potential, and heretofore, the occurrence of action potential.

### B-II The engram mechanism

In Figure 6c, at initiation of AP, *Na*^+^ enter the axon, *K*^+^ is exiting, the amount of *K*^+^ that cross the cell membrane during a single action potential is extremely small relative to the total concentration of inside the cell. The molar amount is estimated approximately to 3.3 × 10^-18^ moles of *K*^+^, per each action potential on a single axon and negligible. The amount of *Na*^+^ crossing the membrane during a single action potential is also relatively negligible compared to the total intracellular concentration, allowing the cell to maintain gradients without constant, immediate correction. That is 0.8 to 1.5 picomole/cm^2^ (or 0.8 – 1.5 10^-12^ moles) of *Na*^+^ cytoplasmic membrane area of the neuron. All of which cannot be considered as an homeostatic disruption.

The inference, and likely the only one to store the information, and engram the LTP, is as follows. Instead of returning intra-cellular within the axon, during repolarization, is to trap *K*^+^ ions into the multiple superposed myelin layers in the physical process as described in Fig. 6 and Fig. 2c (charge trap).

When the action potential propagates along the axon, positive charges (*Na*^+^) enter the neuron. This is equivalent to an electromagnetic field creating negatively charges outside the axon toward the outward concentric multiple myelin layers, acting as charge trap zone^53^. In this case, the myelin layers behave as a dielectric barrier and the applied electric field is strong enough, the charges extend across the Ranvier nodes, creating a non-negligible probability that the negative charges will appear on the opposite side within the multiple myelin layers (charge trap)^53-56^. This process, known as quantum tunneling^54-56^, infer that ions *K*^+^ may remain trapped in a certain architectural organization.

An image is stored in the charge trap flash (CTF) of a microprocessor by converting its visual data into binary values (0s and 1s) and trapping electrons in specific physical locations within memory cells. Unlike traditional flash that uses conductive floating gates, CTF uses a nonconductive layer to hold electrons in place. In the superposed multiple, concentric, spiraled, and highly compacted myelin wraps specifically assembled in 3D may allow such storage of charges by the disjointed 3D spatial superposition (Fig. 6a, 6c). In this case, stored charges are positive ions, with similar effects whether charges are negative or positive charges. The electromagnetic field will be created all the same. To “write” data, following an action potential, this voltage applies to the control gates of the myelin layers producing a ionic charge injection. This causes charges to gain energy and “tunnel” through the myelin layer across Ranvier nodes, via tunneling^54-56^, and deep into the concentric myelin multilayers (Fig. 6b). This is creating an insulated trapping of *K*^+^ ions isolated from the system. Because myelin layers are an insulator, the injected ionic charges (*K*^+^) become physically stuck in “traps” within the myelin lipid (lipoprotein) structure. The *K*^+^ ions may not move freely, completely trapped after the AP (electromagnetic field) ceases. Mirroring a single-bit, a single-level-cell, SLC, a trapped ionic charge (ions *K*^+^) may represent a “1,” while an empty cell (absence of *K*^+^) represents a “0”. The presence of trapped electrons increases the threshold voltage of the transistor; a similar phenomenon may occur in the brain for the ions *K*^+^. This brain apparatus may also behave as a current multi-bit, MLC (2 bits/cell), TLC (3 bits/cell), or QLC (4 bits/cell), such as the modern 2026 3D NAND SSD, solid-state drive to stack large number of non-volatile flash memory cells (NAND). All things considered, the brain may store multiple bits per “cell”, or minute-masses of cytoplasm trapped between fused layers of cell membranes, virtual spaces of relative “negative or lower pressure points” likely important for adhesion and trapping by precisely controlling the number of *K*^+^ ions trapped creating different voltage levels (*e*.*g*., 8 levels for 3 bits). Hereby, the trapping of *K*^+^ ions would be proportional to the AP intensity. All of which would be in line with neuronal plasticity, synaptic strengthening, after repeated use and reinforcement learning^44,47,95,96,116,117,119,122.^

If the homeostatic balance of *Na*^+^ and *K*^+^ cannot be maintained in some ICU patients with life-threatening cardiorespiratory and renal failures, a normal brain function cannot be sustained and coma may result. Electrolyte abnormalities and ensuing failing action potential is one of the causes of brain dysfunction. High levels of extra-cellular *K*^+^ concentration may be associated to a neuronal firing pattern, mixed-mode bursting, MMB^140^, which is never attained in physiological action potential depolarization. When *K*^+^ ions are detrapped from myelin and oligondendria wraps, they may trigger action potentials causing depolarization. Indeed, a slight increase of extra-cellular *K*^+^ ion reduces the concentration gradient across the neuronal making the resting membrane potential less negative, closer to the threshold making it more excitable and prone to propagate AP^141^. A slight rise of 3mmol/L sufficiently the resting membrane potential toward the threshold making it easier to trigger and sustain AP. The Neuropixels 2.0 probe^142^, or rather, the Neuropixels Ultra (NP Ultra)^143^ measuring a single-neuron, single spike resolution, with silicon probe with much smaller and denser to probe extracellular waveform may bring further insight. To support this inference, a resolution in the nano range exploring the minute-masses of cytoplasm trapped between fused layers of cell membranes, virtual spaces of relative “negative or lower pressure points” likely important for adhesion and trapping the *K*^+^ ions would be needed in a complex experimental design. Studies found a mechanical displacement of the axonal membrane accompanying the electromagnetic potential^82^. This mechanical displacement potentially exposes the charge trap (Figs. 6c and 7) through the Ranvier nodes across the myelin-dielectric via a tunneling process. The potential energy is stored in elastic properties of the neuronal membrane and cytoskeleton while kinetic energy is carried by the axoplasmic fluid. These surface waves are driven by the travelling wave of the AP-electromagnetic depolarization, modifying compressive electrostatic forces across the membrane^82^. This potential difference is also propagating mechanical displacements, which the authors called “action waves”^143^. We also infer that this geometrical deformation may facilitate the tunneling and also that *K*^+^ ions become trapped in minute-masses of cytoplasm, joint surfaces forcing apart (tribonucleation), between fused layers of cell membranes, exposing virtual spaces of relative “negative or lower pressure points” to let the *K*^+^ ions enter and being trapped, then the “action wave” clips them in. Sequestering those *K*^+^ ions may be indefinite, until a new sufficiently AP stimulus above threshold (*e*.*g*., a memory recall) reopens those “dormant myelin-memory cells”, further detrapping *K*^+^ ions, generating intense AP.

**Fig. 7.**
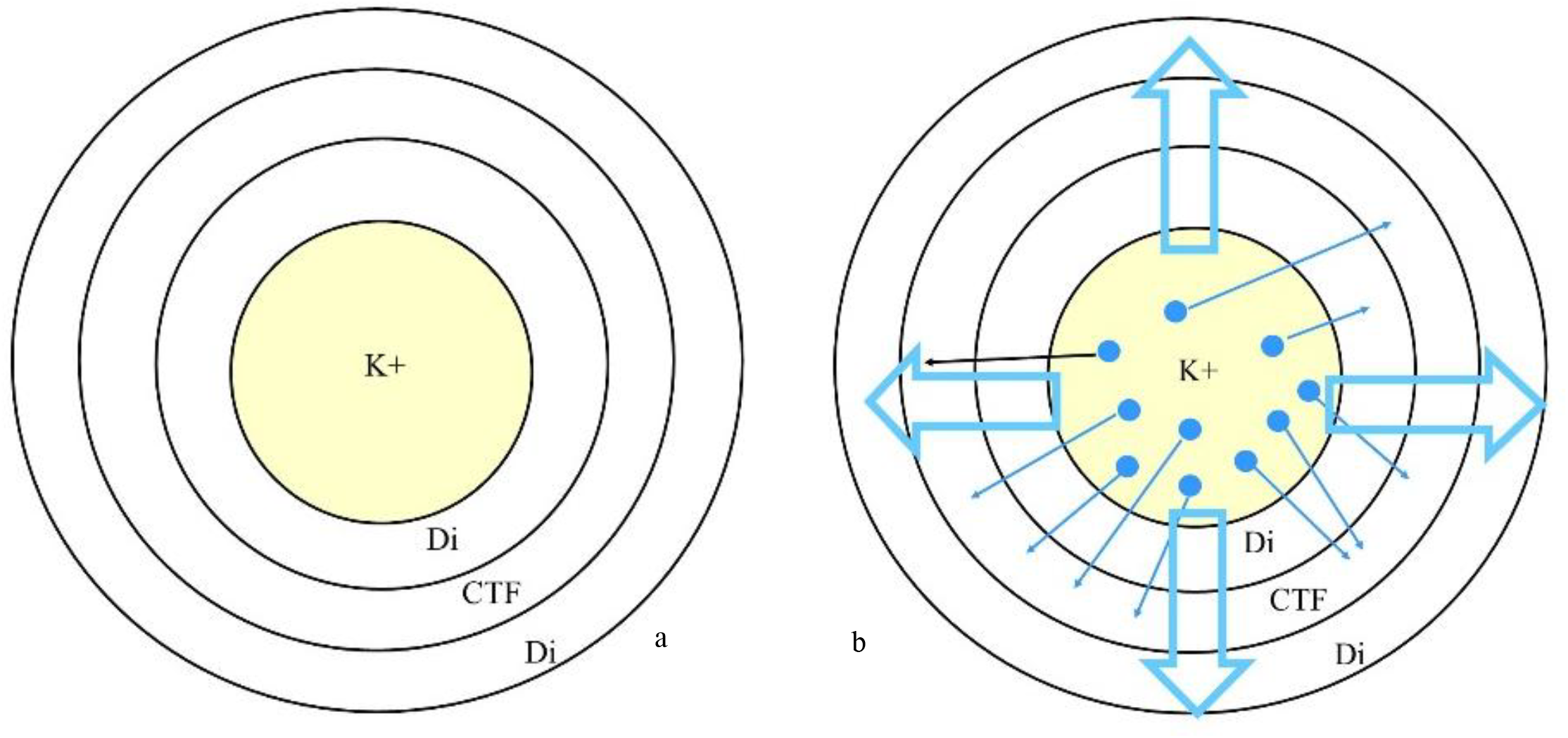
a). The axon and its concentric myelin multilayers restricted to two layers and a virtual space, “CTF”, charge trap flash, fused cell oligodendrocyte membranes. Di, dielectric, is a single myelin wrap. b). the action potential creates an electromagnetic field, then ion *K+* migrates in the extracellular milieu, and across the Ranvier nodes further into the virtual space between two myelin-dielectric layers. Some *K+* remaining trapped, and non-exposed, are storing nonvolatile memory in a CTF-like. The occurrence of a minimal action potential will lower the threshold and liberate the *K+* ions (as memory recall or stimulation). The data may be stored in 0 or 1. See text for details. Large transparent arrows with blue outline are the electromagnetic potential generated by AP. The trapping may occur in a myelin layer of the axon within or rather in adjacent myelin layers compressed in a bundle. See transversal section on Fig. 8 for details.

## C. Discussion

### C-I. Quantum coherence of the isolated system of myelin wrap-ion channel *K*^**+**^**/*Na***^**+**^

#### 1. Quantum coherence – decoherence

Influential physicists, including Penrose^144-146^, Hawking^144^, Wigner and others^147-150^ devised on consciousness, brain, and quantum mechanics. While consciousness is not the focus of this work, considerations of brain function and quantum physical principles are relevant. Quantum decoherence refers to the process where fragile quantum systems lose their unique quantum properties (such as superposition) and default to classical behavior due to interactions with their surroundings (wet/dry, “heat bath”). It was argued that interaction with the environment is probably small enough to be unimportant for certain neural processes^147^ whereas others have proposed that large-scale (“macrosuperposition”) coherence in the brain is unlikely due to rapid decoherence^148-150^. Interpretations of quantum mechanics also depend on environmental context, as system–environment interactions may contribute to effective wave-function collapse. “Dry” environments refer to isolated, low-temperature, solid-state or vacuum systems such as those used in quantum computing. In contrast, “wet” environments refer to thermally noisy, aqueous biological systems such as cells and proteins. While heat generally destroys quantum coherence, recent advancements in quantum physics and biology demonstrate that coherence can be preserved or dynamically maintained even in warm environments, and at or near physiological temperatures (*e*.*g*., 37°C)^151^. Although at 37°C, particles have a high amount of thermal energy, causing rapid vibrations typically collapsing the wave function. Coherence can persist if the quantum system is isolated from its environment such as trapped in a highly rigid molecular structure such as myelin wraps. The environment dynamically re-coheres the system faster than it decays if the observer’s measurement timeframes are extremely short (e.g., femtoseconds or nanoseconds)^151^.

Recent advances in neuroscience and quantum mechanics have significantly expanded our knowledge and may help elucidate the underlying mechanisms operating at the interface between physiology and physics.

#### 2. Specific anatomy-histology preserving quantum coherence

The node of Ranvier is populated by voltage-gated *Na*^+^ channels, *Nav*, adjacent lies the paranode where the myelin is attached to the axon, next to the juxtaparanode where most voltage-gated *K*^+^ channels are located^152^. Voltage-gated *Na*^+^, *Nav*, channels, concentrated at the nodes, are separated from *K*^+^, *Kv1*, channels, clustered at the juxtaparanodal region by a specialized axooligodendroglial adhesion-contact that is formed between the axon and the myelinating cell at the paranode^153,154^. Note that such *Nav* vs. *Kv1* separative property avoids potential decoherence by charged ions collisions. The paranode acts as a diffusion barrier which separates the *Nav* channels in the node of Ranvier from the *Kv1* channels in the juxtaparanodes (Fig. 8). The paranode “glues” the myelin to the axon, ensuring a tight connection between the myelin and the axolemma, and reducing current flow under the myelin sheath^152^. The strong adhesion between myelin and axolemma is achieved by cell adhesion molecules (see below)^152^. The juxtaparanode, adjacent to the paranode, where the axon is directly under compact myelin wraps, but does not extend for the entire length of the internode. Juxtaparanodes are clustered by Shaker-type *Kv1* channels.

**Figure 8.**
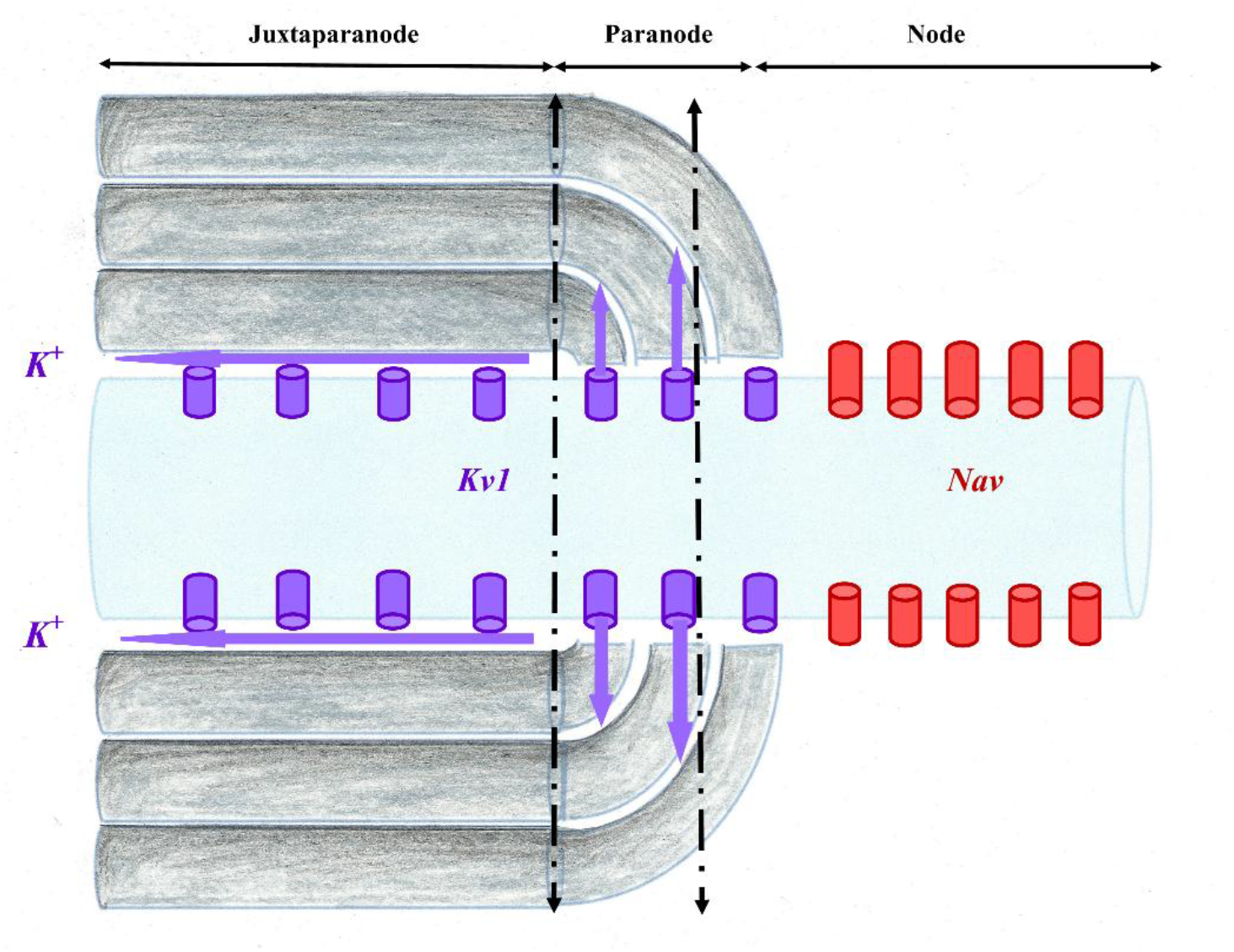
Node, paranode and juxtaparanode. *Nav* ion channels in red. *Kv1* ion-channels in purple. Ions *K*^+^ may circulate within the axooligodendrial space (purple arrow) on the external surface of the myelin wrap. Storage of ion *K*^+^ occur externally and with two myelin wraps. Within a bundle, compressed myelin wraps are extremely hydrophobic, repelling water molecules and perfect insulators. From the outside in, the layers of myelin membrane compact, are bringing the inner and outer leaflets of consecutive lipid bilayers in extremely close apposition and extruding cytoplasm and extracellular fluid. In quantum mechanics, the space in-between can potentially be considered as within a topological insulator. This unique setting is likely preserving quantum coherence of the system. Once quantum tunnelling occurred, ions *K*^+^ might be stored in the space between two myelin wraps. The overall system becoming potential CFT memory cell. Dotted lines: transversal sections are also expanded on a *2-D* plane in Fig. 7.

The high density of *Nav* channels generates the maximum transmembrane voltage across the nodal membrane^155^. Any *Nav* channel misplaced to the internodal region wrapped with multiple myelin layers of myelin wraps, would become dysfunctional and generate a smaller voltage change. Most voltage-gated *K*^+^ channels, *Kv1*-type, are inserted in the juxtaparanodal region^152^. *K*^+^ channels, *Kv1* do not normally participate in generating AP. Blocking or knocking out *Kv1* channels did not affect propagation of PA^152^. Potential current flow is reduced by adhesive junctions between the ensheathing glial cells and axons^152^. These septate-type tight junctions leave a virtual space to the spiral pathway between the extracellular space at the node and the extracellular space at the juxtaparanode, through which some difference of potential (*K*^+^) may expand allowing partial activation of juxtaparanodal *K*^+^ channels. The exact level of AP activation depends on the current flow occurring under the myelin at the juxtaparanodal region, to maintain the extra-axonal voltage close to 0 mV. Structures associated with *Kv1* channels are found in the internodal and paranodal regions and are normally concealed under the myelin sheath^156^. The acceleration of myelin swelling after repetitive propagation of APs may be due to the opening-loosening of paranodal axooligodendroglial junctions^156^. From the outermost to the innermost layers, the myelin membrane undergoes progressive compaction, resulting in the close apposition of the inner tongue and outer leaflets of adjacent lipid bilayers. This process is extruding both cytoplasmic and extracellular fluid, thereby generating the highly compact structure characteristic of mature myelin^157^.

Nodal length and diameter are strictly regulated. The speed of nodal *Nav* channels activation depends on the capacitance of the node, and hence by its membrane area^152^. Pathologies increasing the area are slowing AP, unless extra *Nav* channels are inserted into the membrane to oppose this effect. Furthermore, an increase in node length, as also occurs in pathology, will increase the axial resistance for the current flow from the node to the internode that is needed to depolarize the next node, again slowing the action potential, or even causing it to fail^152^. Pathological disruption and retraction of the paranodal structure may loosen this junction^158^, ensuing preventing AP propagation, especially during repeated firing. Loss of internodal and paranodal adhesion results in overgrowth of nodes of Ranvier^159^. In multiple sclerosis, cerebral hypoperfusion and energy deprivation lead to elongation of the node of Ranvier, ensuing *Nav* channels covered with myelin wrap and disruption of paranodal junction, by loss of cell adhesion molecules (neurofascin 155, NF155), and paranodal contactin-associated protein (Caspr)^152^. This increases the virtual axooligodendroglial space between axolemma and myelin wrap, and *Nav* channels overlap with the paranodal protein Caspr^152^.

In addition, potassium channels *Kv1* within the juxtaparanodal axonal membrane may serve as depressing re-entrant excitation^160,161^. The astrocytic processes contain the inwardly rectifying *K+* channels *IRK1* and *IRK3*, which may allow *K*^+^ to accumulate during axonal activity^160,161.^

#### 3. Axooligodendroglial space and quantum coherence

The extracellular perinodal matrix is not a fluid or a liquid. Oligodendrocytes act as the primary producers of this specialized ECM medium. It takes the form of a dense, localized, gel-like macromolecular network that is structurally unique from the rest of the brain’s general interstitial matrix^162^. The molecular composition of the perinodal medium, rather than being fluid, liquid or generic connective tissue, this local medium is densely rich in specific structural and signaling macromolecules such as chondroitin sulfate proteoglycans, CSPG, highly negatively charged, hydration-retaining physical backbone of the node; brain link protein 1 anchors the proteoglycans into stable, higher-order aggregates; glycoproteins interacts directly with the axonal surface; hyaluronan the universal structural scaffold binding the proteoglycans^153,155.^ The role of the perinodal matrix is ion buffering via the intense negative charge of the CSPGs creates a localized cation reservoir. This helps manage the rapid influx and efflux of *Na*^+^ and *K*^+^ ions required during saltatory electrical conduction^155,163^. It is also to molecular anchoring, It physically stabilizes and secures the clusters of voltage-gated sodium channels Nav on the exposed axon, keeping them from drifting into the myelinated segments^163^.

Axial conduction through the axooligodendroglial space is determining the saltatory AP propagation of action^157,164^. Modulation of the axooligodendroglial space provides degrees of structural plasticity to the system^164^. Spatial learning alter the size of the axooligodendroglial space^164^. Modulating neuronal activity induces adaptive changes in myelin ultrastructure altering length of the node of Ranvier and width of the axooligodendroglial space (normal ≈ 12 nm), two of the axoglial parameters strongly determining AP conduction velocity, *V*. Furthermore, the inclusion or omission of axooligodendroglial space width could result in new internodes increasing or decreasing *V*, respectively^164^. Electrical conductance within the axooligodendroglial space facilitates the saltatory propagation of AP and changing the volume of this space theoretically alter *V*^164^. This property has implications for quantum tunnelling of brain memory cells. Simulated *V* up to 3.5 times slower when the axooligodendroglial space width is set to 20 nm rather than 0 nm^164^.

#### 4. The unique system myelin wrap-ion channel *K*^+^/*Na*^+^ isolating from microscopic environmental disturbances

Working memory training stimulates the production of myelinating oligodendrocytes from their precursor cells in the medial prefrontal cortex, anterior corpus callosum, dorsal thalamus and hippocampal formation^165^. Myelin sheath formation is also stimulated in the genu^165^. Myelin (white matter) is a highly hydrophobic structure composed of approximately 80% lipids, including galactocerebrosides and long-chain sphingomyelins, whose densely packed hydrocarbon chains confer strong water-repellent and electrical insulating properties. The remaining ∼20% consists primarily of structural proteins, proteolipid protein (PLP), among the most hydrophobic proteins known, whose transmembrane helices are deeply embedded within the lipid-rich membrane core. In the brain, myelin is proportionally thicker in thinner axons, measured as a *g*-ratio by electron microscopy^157^. The memory cells embedded within concentric 300 myelin layers wraps can be considered as dry solid-state relative vacuum (Fig. 8). Those virtual anatomical spaces of relative “negative or lower pressure points” are important for trapping in isolated niches of dry environment. Most biological mediums are made of fluids; however, specific anatomical 3D-organization of myelin wraps introduces localized conditions that are providing a dry, solid-state or vacuum-like conditions (relatively low-pressure environment) mitigating decoherence effects. This anatomical brain specificity makes it dry-hydrophobic with low-pressured air vacuum. The anatomical particularity of myelin wraps is that the *2-D* plane of unfolded myelin layers exposing the random disjointed placement of Ranvier nodes exponentially increasing the hosting of charges ions in their memory cells. This is also protecting the system myelin wrap-ion channel *K*^+^/*Na*^+^ from the fluidic environment and from firing, AP, because there are ∼300 myelin wraps around the axon in a bundle that may contain up to 200-200 million compress myelinated axons going from *2-D* to *3-D* or to a Hilbert Space. All of which, disjointing and isolating Ranvier nodes, only allowing very limited localization of transfers and AP propagation; this also warrants the precision of the signal in a bundle. Axons within a single long white matter bundle fire asynchronously at different times. Figure 6 illustrates the translation *3-D* to *2-D* allowing this asynchronous property and near-perfect isolation of the myelin wrap-ion channel *K*^+^/*Na*^+^ system allowing quantum coherence. It is randomly unlikely that two Ranvier nodes from two axons are facing each other. Hereby, given the myelin hydrophobic isolating the firing, rapid decoherence timescales 10^-13^ - 10^-20^ s within fluids will not affect firing timescales 10^-3^ - 10^-1^ s. Given the highly hydrophobic myelin wraps, collisions with water molecules is unlikely in the isolated myelin wrap-ion channel *K*^+^/*Na*^+^. In a bundle, myelin wraps are compressed and the oligodendrocyte intracellular space within the myelin wrap is likely virtual and squeezed enabling perfect insulation. Collisions with other ions is possible although unlikely since the general ion motion is mainly one-direction and because the cloud motion *K*^+^/*Na*^+^ in opposite direction occur at subsequent timeframes and in different locations *K*^+^ in paranode and juxtaparanode, *Na*^+^ in the node.

#### 5. Unique physical properties conferred by the node, paranode, and juxtaparanode

Ion channels *K*^+^/*Na*^+^ pumping (ATP-energy-system) driving ions against their electrochemical gradients, ***moving ionic charges, Na***^**+**^, ***K***^**+**^, in nodes, paranodes and juxtaparanodes, are creating an ***electromagnetic, EM, field***. It expresses a voltage, nodal intra- and extra-neuronal potential difference, which can be measured^95,166,167^, although action potential along the axon is ***not*** a conventional electric current relying on physical flowing of electrons through a conductor. The presence of the myelin sheath insulator, extracellular matrix and cells create a unique environment for propagating this EM field. Each ion channel *K*^+^/*Na*^+^ creates self-propagating electromagnetic waves jumping to the next ion channel, triggering the next wave, from next channel to next. This unique setting of myelin wrap-ion channel *K*^+^/*Na*^+^ systems, energy-dependent at each ion-channel, is an overall total energy-saving. Electromagnetic waves travel at the speed of light (299.10^6^ m/s) in perfect vacuum which is not the case across the extracellular matrix, or cytoplasm dropping the velocity down to 194.10^6^ to 214.10^6^ m/s. This EM wave velocity might be dropping through insulators like myelin sheath down to 124.10^6^-150.10^6^ m/s. This is approximately the propagation velocity of EM from node to node and start triggering the activation process of ion channels *K*^+^/*Na*^+^ pumping at the next node. Indeed, this is not the propagation velocity of AP, *V* ≈ 48 – 150 m/s. This drop of *V*, AP velocity, is time for ATP energy recruitment and activate the ion channels *K*^+^/*Na*^+^ pumping. The myelin sheath serves as an insulator for charged ions, from one node to next and as a dielectric in a node, paranode and juxtaparanode. Myelin might also behave as topological insulator for the EM waves, electrical insulator inside, but conduct electricity perfectly along the myelin-wrap surface, across extracellular matrix, or cytoplasm (Fig. 8). An EM wave triggers a new phase in distant charged ions primarily through mechanisms of EM-driven phase transitions^168,169^ such as for instance, by resonant energy transfer^170,171^ with potential phonon-phonon coupling^172,173^ assuming a myelin-wrap may behave as topological insulator (*i*.*e*. “quantum material”), or by field-induced polarization^170,174^. When the oscillating electric field of an EM wave matches the natural frequency of a targeted ionic lattice or plasma, it drives collective structural transitions even across significant distances. Low-energy cyclotron emission is an established physical phenomenon. It occurs whenever any charged particle (like an electron) spirals in a magnetic field at low or non-relativistic energies^175^. The particle emits electromagnetic radiation at the cyclotron frequency, which is directly proportional to the magnetic field strength and inversely proportional to the particle’s mass.

Direct visualization of fast APs propagating along internodes remains infeasible due to EM propagation over a short internode length^176^. The dynamic phase-event takes several femtoseconds to propagate through a micrometer-scale internode; hence, it is too fast to be captured using existing phase imaging techniques relying primarily on binning temporal frames to improve sensitivity^176^. EM can propagate near the speed of light depending on the dielectric media^176^, such as myelin (124.10^6^-150.10^6^ m/s). To noninvasively, flexibly image the propagation of the fast transient excitation frame rates of up to 200 billion frames/s, a sequence depth of 350 frames, and the high phase sensitivity, differentially enhanced compressed ultrafast photography, Diff-CUP achieves noninvasive, direct spatiotemporal observation of passive internodal flows of electrical current, which plays a central role in AP propagation, and transmission of any other forms of electrical signaling across internodes where ion channel activities are absent. With a frame rate of up to 200 billion fps, a sequence depth of up to 350 frames, and a phase sensitivity of 20 μrad, Diff-CUP has enabled direct imaging of internodal current flows propagating in myelinated axons with an average conduction speed of 100m/s as well as the direct visualization of an EMP propagating in an LN crystal with a speed of 5 × 10^7^m/s. The reproduced internodal current flow speed and LN relative permittivity are consistent with the simulated and reported values, demonstrating the translation of our method into the biological and physical sciences. This team has reported compressed ultrafast photography (CUP), the world’s fastest camera that achieves both an ultrafast imaging of up to 70 trillion frames per second (fps) and a large sequence depth of up to 1000 frames^176^. However, without sufficient phase sensitivity (*e*.*g*., 0.9 mrad required to measure Aps) in spiking their previous CUP systems could not image either propagating APs or weak EM Pulses^176^. It is likely that an accurate assessment of the quantum physics underlying AP in the human brain will soon be made.

### C-II. The special case of Alzheimer’s disease provides further evidence for the proposed mechanism of memory storage

Oligodendrocyte and myelin disruptions are paramount factors in Alzheimer’s disease, AD, although the mechanisms remain unknown. Characteristic amyloid deposits, including spiral fibers around axons and dense helical coils at paranodal-juxtaparanodal loops, indicate amyloid infiltration within the axooligodendroglial space at paranodes, juxtaparanodes and internodes^177^. Additionally, we found aberrant myelination and disrupted paranodes on amyloid-plaque-associated dystrophic axons, a hallmark of AD known to impair action potential conduction and disrupt neuronal networks^177^. Despite no significant difference in the overall myelin coverage, there was greater localised paranode loss in AD brains compared with age-matched controls, indicating dysregulation of myelin-related signaling rather than a net loss of myelin. These results highlight significant paranodal protein alterations in AD, underscoring molecular disruptions at the axooligodendroglial space that may contribute to this pathogenesis^177^. Paranodal and periaxonal myelin regions are sites of amyloid-β accumulation^177^. This adds thickness and insulation within the axooligodendroglial space at paranodes, and juxtaparanodes, further blocking and preventing ions *K*^+^ to cross the axooligodendroglial space at paranodes, and juxtaparanodes. This would suggest that such thicker myelin wrap cannot serve as thin dielectric to achieve quantum tunnelling and *K*^+^ cross out to memory cells within myelin wraps. Amyloid-β processing is dysregulated in the axooligodendroglial interface proteome and dense amyloid coils originated at paranodes and extended into juxtaparanodes^177^. The dense helical patterns likely represent amyloid-β accumulation within paranodal channels, while the less dense spirals may be extensions into periaxonal spaces at internodes. Amyloid spiral and coil around an axon within the axooligodendroglial space are localized at paranodes and juxtaparanodes^177^. This invasive axooligodendroglial pathological process may prevent normal quantum tunnelling into thicker dielectric myelin layer. Bulbous axonal enlargements were sometimes observed at these helical amyloid-β aggregation sites. All of which occur without overt myelin loss^177^, rendering difficult the imaging diagnosis in AD patients.

### C-III. Conclusion

The human brain design favored a packed, multi-function circuitry, to optimize the volume, weight, energy, efficiency, rapidity, precision, and storage. The same conductive input circuitry is reused in memory cells and vice versa according to the selective inhibition or reinforcement. The human brain model was selected over an exhaustive infinite accumulation of data, depending on immense energy-consumption, heat production, weight and volume beyond the capacity of the human skeleton/muscles architecture to freely and easily move in one-gravity.

## D Methods

Analyzing the complex, multidisciplinary mechanisms of brain action potentials requires careful categorization and rigorous examination of a broad range of interconnected factors, *including relationships and interactions that may not be immediately apparent*. In the present study, we focus on recent biometric data, and distinct velocities of action potential and electromagnetic-driven phase transitions. Rather than evaluating competing theoretical paradigms and imaging, our analysis is restricted to quantifiable metrics, specifically propagation velocity and energetic cost. To investigate these relationships, we performed computational statistical modeling of the recent biometric data using Monte Carlo simulations implemented in *R*.

Our sensory systems may collect data at ∼10^9^ bits/s^111^ enabling remarkably fast reaction times. A tennis player can be judging the speed (∼ 130 MPH), the strength, future geo-localization of the ball in space, forecasting its direction, running, dodging, anticipating the body position, racket placement, optimizing the shoulder-arm lever in space, while briefly checking the angle on the other side of the net, and hitting the ball with slice effect for a winning point. All of which, accomplished in a fraction of second.

In comparison, the processing speed of the brain, information throughput of a human brain was evaluated ∼10 bits/s, relatively small^111^. This was evaluated based on ∼1 bit/character in English language, the brain seems to produce an output of ∼10 characters/s or ∼10 bits/s. In this supplementary information, we are interested in the relationship between the *qubit*-information broadcast, the energy/*qubit* ratio, and the comparative speed of transmission/processing of information. Recent biometrics of the visual pathway^178^ and other studies^63,179-181^ enabled to determining speeds and comparing respective times of processes to provide further insights into the roles of brain structures. We are investigating this intriguing duality of a low energy-fast-brain and high energy-slow-brain based on timing and biometrics.

### D-I. Recent human brain biometrics data

To account for variations across individuals, we ran Monte Carlo simulations on the lengths of each segment as provided by Pravata, E. et al.^178^ for the anterior visual pathway (Segment 1, 2, 3, 4, & 5) and from Peltier, J. et al.^182^ for the posterior segment. The six segments were Orb, intraorbital segment; Can, intracanalicular segment; Cran, intracranial segment; OC, optic chiasm; OT, optic tract; and posterior visual pathway (Fig. 9, adapted from Takemura et al.^179^). The authors^178^ found no statistically significant difference between the left and right side for each parameter (*p-value* range 0.249–0.931) and for Segment 6 only one value is given^182^. We used and only reported the data relevant to our research (Table 1)^178^. *Positive values indicate left > right, and vice-versa. In our program, we included the time to transfer the *qubits*/electron spin – spike, through the cleft via *qubit*/ion transfer or other, to post-synaptic back to *qubit*/electron spin – spike. The biologist’s approximation, “one bit in one neuron spike”^63^, may be considered as a mesoscopic raw extension to several *qubit* – electron spins necessary to make one bit of a spike. The crossing time through the intersynaptic cleft, reported by Markram et al. was 1.7 ± 0.9 ms^166^, for which we also ran Monte Carlo simulations is the seventh step as referred in *Eq. 1*. At this point, we had not yet made the relation with quantum physics; we were still using the terminology for electric current with electrons ignoring the transversal transfer across cellular membrane: ion channels *K+/Na+* pumping (ATP-energy-system) driving ions against their electrochemical gradients (see text).

**Table 1.**
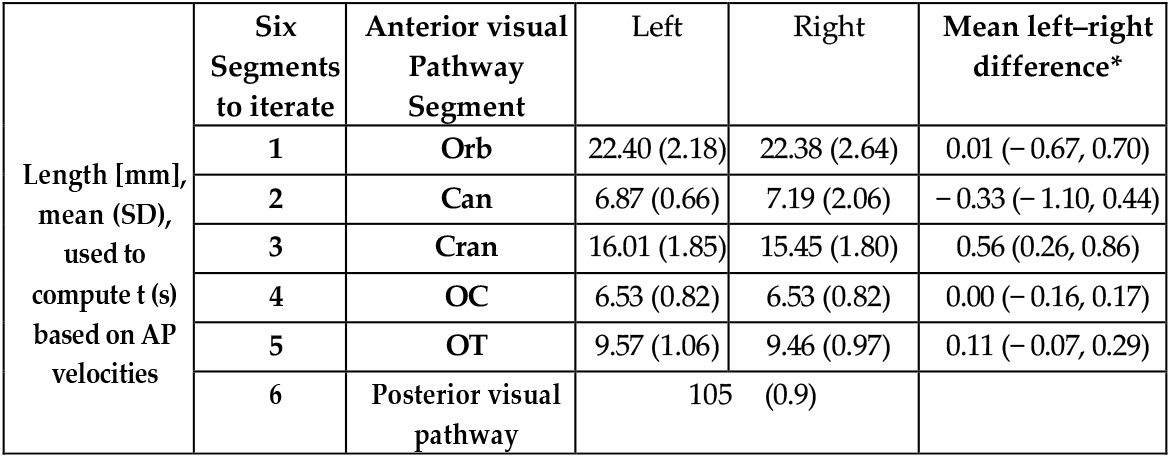
Visual pathway. Segment lengths (mm).

|  | Six Segments to iterate | Anterior visual Pathway Segment | Left | Right | Mean left-right difference* |
| --- | --- | --- | --- | --- | --- |
| Length [mm], mean (SD), used to compute $t$ (s) based on AP velocities | 1 | Orb | 22.40 (2.18) | 22.38 (2.64) | 0.01 (-0.67, 0.70) |
|  | 2 | Can | 6.87 (0.66) | 7.19 (2.06) | -0.33 (-1.10, 0.44) |
|  | 3 | Cran | 16.01 (1.85) | 15.45 (1.80) | 0.56 (0.26, 0.86) |
|  | 4 | OC | 6.53 (0.82) | 6.53 (0.82) | 0.00 (-0.16, 0.17) |
|  | 5 | OT | 9.57 (1.06) | 9.46 (0.97) | 0.11 (-0.07, 0.29) |
|  | 6 | Posterior visual pathway | 105 (0.9) |  |  |

**Table 2.** These results are merely in line with a thorough literature search across hundreds of papers. Values are means and SD in seconds. From West et al.

| HRF parameter | Young | Old | P |
| --- | --- | --- | --- |
| Time-to-Peak (s) | 4.194<br>(0.083) | 4.652<br>(0.096) | 0.003 |

**Figure 9.**
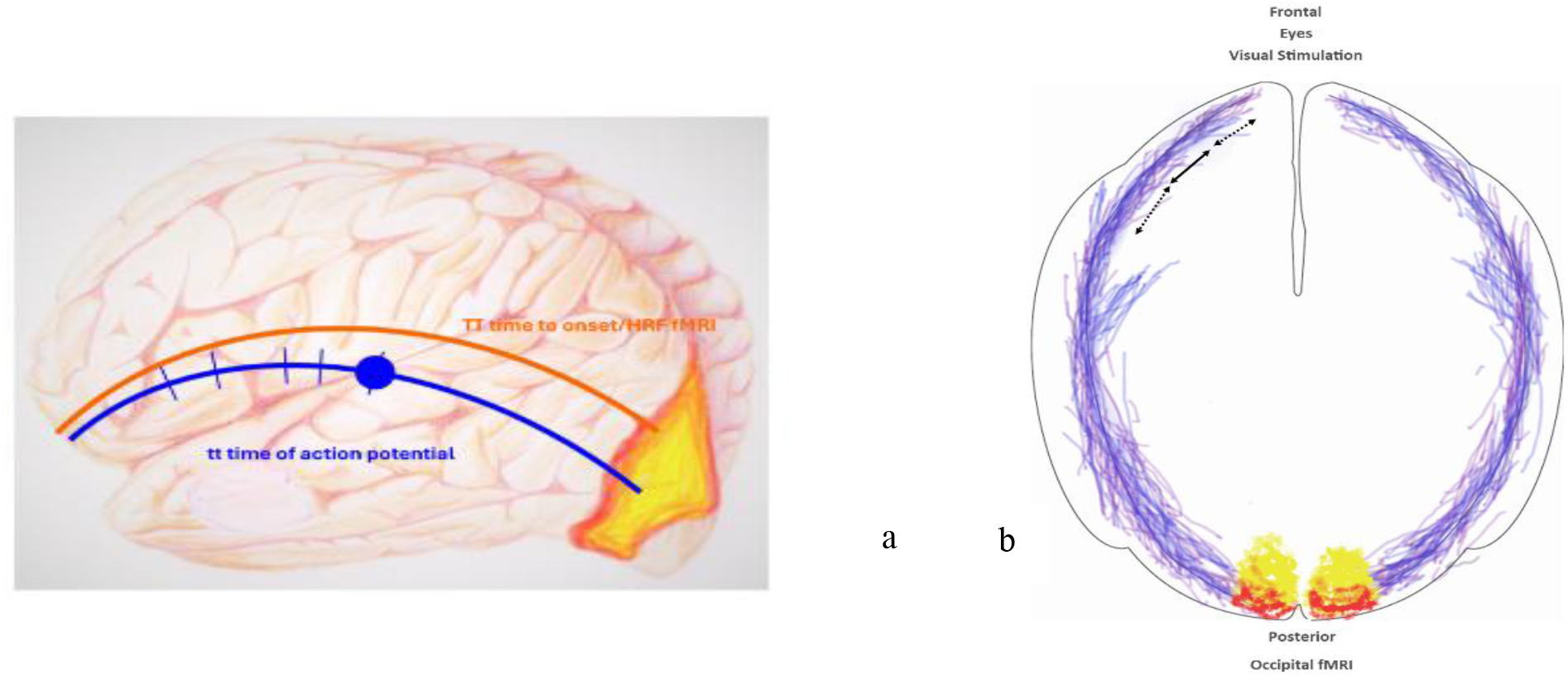
**a**). Simplified non-anatomical representation of the action potential trajectory from the intra-orbital inception of the optic nerve to the occipital cortex, region of interest, ROI, via the visual pathway. (optic chiasm is not represented). In blue, the action potential. The ticks are an unscaled representation of the segment lengths that we study. The blue circle is one lateral geniculate nucleus, LGN. In orange, we illustrate the simplified trajectory of the action potential which will generate the BOLD Hemodynamic Response Function, HRF, and directly visualized as voxels on the fMRI. **b**). Simplified and non-anatomical representation of the visual pathway. Ipsilateral and contralateral pathways, optic chiasm and lateral geniculate nucleus (LGN) are not represented. Ipsilateral and contralateral does not affect our mathematical computation. visual areas showing maximal blood flow (activity) during visual pattern stimulation a. hottest voxels in red. b. The double-arrow illustrates the measurement of anatomical segment lengths.

### D-II. Functional MRI data

The BOLD Hemodynamic Response Function, HRF, was assessed by West et al.^180^ in two samples of 74 younger participants (18– 30 years of age; SD = 3.58) and 173 older participants (54–74 years of age; SD = 6.00)^180^. The authors collected the magnetic resonance imaging (MRI) as part of a one-hour session in a 3T Siemens TIM Trio scanner with a 32-channel head-coil (Fig. 9)^180^.

Sample mean HRFs were different between younger and older groups in visual cortex. The authors observed increased time-to-peak and decreased peak amplitude in older compared to younger adults^180^.

### D-III. Computational statistical modeling

A conversion of consistent units (ms/s, mm/m) was required. The computations and statistical analyses were performed on R Program Version 4.5.0 Alpha (2025).

The action potential, AP, travels from the inception of the optical nerve, Orb, through all segments, Can, Cran, OC, OT, and posterior visual pathway to its final destination, the visual processing occipital area. The time for the AP to cross the intrasynaptic cleft at the LGN levels, was also added as a seventh step. The total travel time (in s) was provided by the Monte Carlo simulations which were computed for each of the seven steps; a total of six segments and the intrasynaptic cleft, LGN (*k* = 1, …, 7). For 10^5^ Monte Carlo simulations, the total travel time 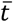 is expressed as,

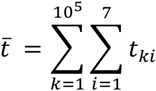

the variance is given by the following formula,

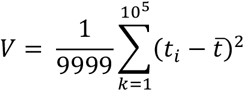

with standard deviation, *SD*, as

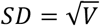

To further calculate the variance and standard deviation of an iterative summation of independent random variables, we used the “aggregates matrix” referring to the organized collection of data for these variables, with each row representing a variable and including its properties, such as its mean and standard deviation such as,

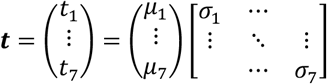

where *μ*_*i*_ are the means and *σ*_*i*_ standard deviations each of six segments and LGN synapse

### D-IV. Analysis of the human brain biometrics data

We performed 10^5^ Monte Carlo simulations iterating six segments, from Orb, intra-orbital inception of the optic nerve, through the anterior pathway and posterior visual pathway, to the visual areas V_j_ (j = 1, 2, 3, 4, …, n).

The Monte Carlo simulations returned to total length with mean = 166.33 mm and SD = 4.07 mm (Fig. 10).

**Figure 10.**
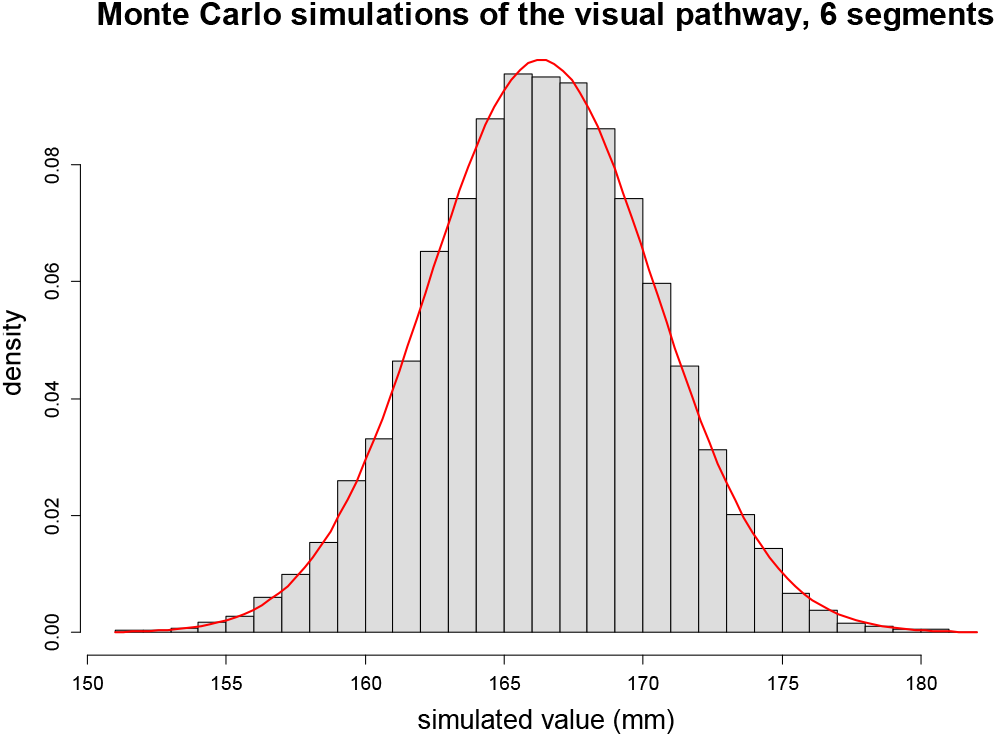
Histogram of Monte Carlo simulations. Iteration of six segments, total length of the visual pathway (mm).

### D-V. Analysis of the velocities

1. For the slowest velocity of the AP (48 m/s), our 10^5^ Monte Carlo simulations for the six segments including the time to cross the intersynaptic cleft (lateral geniculate nucleus) returned a mean = 3.50 ms, and SD = 0.09 ms (Fig. 11).
2. For the fastest velocity of the AP (150 m/s), our 10^5^ Monte Carlo simulations for the six segments including the time to cross the intrasynaptic cleft (LGN) returned a mean = 1.12 ms, and SD = 0.03 ms (Fig. 12).
3. Sample of young subjects, slowest velocity of the AP (48 m/s). We ran a Welch two-independent samples t-test. It returned t-statistic = -5037.7, degrees of freedom, df = 9999, and *p-value* < 2.2 10^-16^. We anticipated that the *p-value* would be highly significant provided the difference in the scale magnitude of t (ms range) and T (s range).
4. Sample of older subjects, slowest velocity of the AP (48 m/s). The Welch two-independent samples t-test returned the t-statistic = -4869.1, df = 9999, and *p-value* < 2.2 10^-16^.
5. Sample of young subjects, fast velocity of the AP (150 m/s). The Welch two-independent samples t-test returned the t-statistic = -5079.7, df = 9999, *p-value* < 2.2 10^-16^.
6. Sample of older subjects, fast velocity (150m/s). The Welch two-independent samples t-test returned the t-statistic = -4871.6, df = 9999, and *p-value* < 2.2 10^-16^.

**Figure 11.**
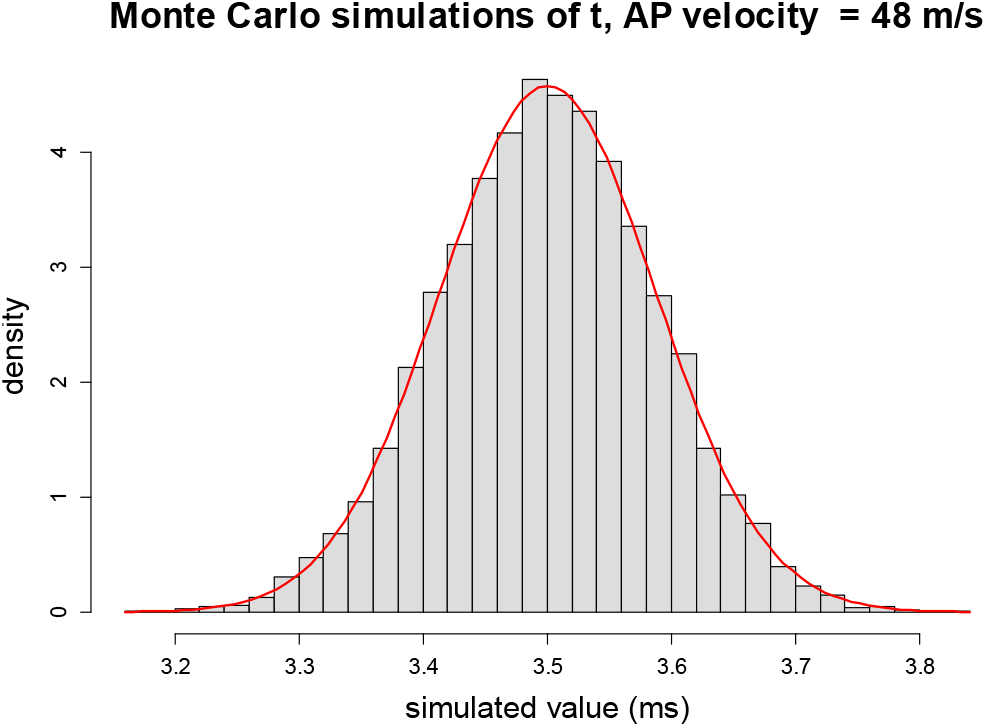
Histogram of Monte Carlo simulations. AP velocity of 48 m/s.

**Figure 12.**
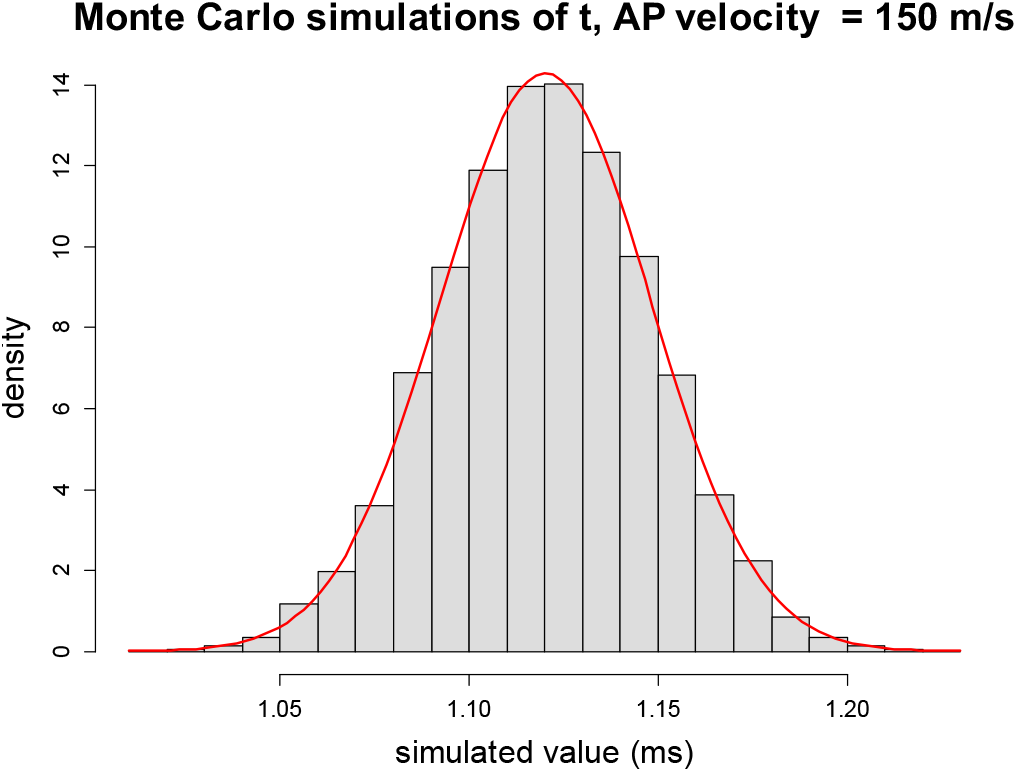
Histogram of Monte Carlo simulations. AP velocity of 150 m/s.

Intuitively, the results were anticipated. The propagation of electricity through a conductor is 150,000-297,000 km/s slightly less than the speed of light. In contrast to the brain, where the action potential is **only** 0.048 to 0.150 km/s, because the remarkable brain wiring is specific, it enables “electromagnetic jumps” over insulation from hole to hole (aka Ranvier nodes) on the myelin sheath; this generates the bouncing and conduction of the action potential. In nodes, where the Schwann myelin sheath is unveiled and the cell membrane exposed, ion pumping creates a potential difference and magnetic field. Indeed, observations constantly showed that difference of scale magnitude between *t* (ms) and *T* (s). However, it was further relevant to check that there was no overlap of the two normal *pdf* distribution at 95%, *t* (+ one SD) and *T* (-one SD) albeit this difference of scale. Indeed, it makes sense to test the null hypothesis for two independent samples from normal distributions with different scale magnitudes^183^; the Welch’s t-test (also called the unequal variances t-test) should be used instead of the standard Student’s t-test, as the latter assumes equal variances and will provide inaccurate results^183^. The null hypothesis would be that the two sample means are equal, and the alternative would be that they are not equal. As anticipated, in all conditions tested, such as age, velocities of action potential, *t*, travel time of the action potential from intra-orbital inception of optic nerve, to the region of interest, occipital cortex, via all segments of the visual pathway, is much different than the time-to-peak of HRF as seen on fMRI.

### D-VI. Velocities of two different scales

Drawing on anatomical data, recent findings on the biometrics of the human brain—particularly the long-range white matter association tracts of the visual pathways—are providing additional insights into the analysis of action potential propagation velocities.

An intriguing duality of a low energy-fast-brain and high energy-slow-brain was based on timing and biometrics. The fMRI, blood oxygenation level-dependent (BOLD) signal, is significantly delayed (*p-value* < 2.2 10^-16^) and appears in the occipital cortex seconds after the AP signal reaches out the visual areas. Such a drop of vascular deoxy-Hb content (fMRI) reflects the ATP-dependent immediate mitochondrion-intracellular utilization of oxygen^49,184-186^. Glucose is considered the main source of energy for neurons and under normal conditions adenosine triphosphate (ATP) is almost exclusively produced by glucose oxidation. The aerobic pathway for high-constant energy production and restoration of a cell’s resting state. Such intracellular process of aerobic respiration requires oxygen in order to create ATP. Hereby, this delayed massive energy for processing information in sub-level dimensions, is likely intracellular, although the specific cells involved are unknown, and may be neurons, astrocytes or vascular cells.

Ability to group visual stimuli into meaningful identified categories of objects is a fundamental cognitive process. Visual stimuli and their delayed match-to-category identification task requires judging. Recordings of 395 neurons from the lateral prefrontal cortex, PFC, of two monkeys led to assessing the reaction times for object identification to be in the range of 2,000-2,500 ms^40^. Across the population of neurons, category activity appeared at the start of neural responses to the sample, about 100 ms after sample onset^40,41^. The hallmark of a classical perceptual categorization is a “classical” perceptual precise category “boundary”. Perceptual categorization depends on extraction of the combinations of features defining a category. These object shapes were not explicitly instructed^40^. They were learned during testing and were necessarily multivariate abstractions; the object categories differed by more than a few simple features. PFC activity could have reflected, and/or resulted in, a shifting of attention to the specific shapes/features of the objects. The PFC is interconnected with temporal lobe structures important for long-term memory, including the inferior temporal cortex, ITC, whose neurons have stimulus specificities that could contribute to categorization^40^. Interactions between the PFC and ITC are the underlying process to recall visual memories and associations, but not necessarily visual short-term memory. The ventromedial PFC through direct projection pathway to the medial entorhinal cortex modulates entorhinal activity, in the dorsal hippocampus (CA1), controls memory integration, allocation and linking^187^.

Remarkably, the brain expresses this dual ability, slow and fast reaction times. A fast conduction tract or “spin” “hopping” from one Ranvier node to the next enabling life-saving quick reaction times. Hereby, this fast response from perceptual, associative integration, and reaction depends on a rapid transmission of information. Based on the extreme velocity, most of which seems to be on white matter tracts.

This led us to re-examine the transversal transfer across cellular membrane: ion channels *K+/Na+* pumping (ATP-energy-system) driving ions against their electrochemical gradients (see paper). This fast conduction system in the brain was not relying on electric current with electrons; the brain’s velocity also was dramatically slower. Furthemore the brain complex fine-tuning equipement for AP, ATP-energy dependent, does not support traditional electron-based electric current. Re-examining the transversal transfer across cellular membrane, ion channels *K+/Na+* pumping, prompted us to investigate existing models of data storage in solid-state drives (SSDs), with particular focus on the underlying mechanisms of quantum physics, including quantum tunneling, charge trap flash (CTF), and dielectric materials. These models offer insights into the interaction of electrical charges and the potential for data retention and retrieval informed by SSD technology.

The propagation of electric signals in conductive materials such as copper occurs at velocities approaching the speed of light, with the effective signal speed, *S*, estimated to range from approximately *S* ≈ 150.10^6^ to 297.10^6^ m/s. Note that the drift velocity of individual electrons within these conductors is markedly slower, on the order of 10^-4^ – 10^-5^ m/s. In contrast to long-distance white matter association tracts of the brain, the velocity, *V*, of electromagnetic field-induced action potentials, is about *V* ≈ 48 – 150 m/s. Although relatively rapid, it is nevertheless much slower than classic electric velocity (*S*) in metallic conductors. Clearly, there is a distinct phenomenon of the electromagnetic field propagation (*V*) occurring in the brain which we explored in the paper.

## Appendix E. R Program. Program repeated for each case (see Fig. 2, 3, 4)

~~~
set.seed(29986577)
library(MASS)
mus <- c(22.40, 6.87, 16.01, 6.53, 9.57, 105, 1.7)
sigmas <- diag(c(2.18, 0.66, 1.85, 0.82, 1.06, 10, 0.9))
ll <- rowSums(mvrnorm(n = 10000, mu = mus, Sigma = sigmas) )
tt1 <- ll/48000
c(mean(tt1), sd(tt1))
### Plot the results
hist(tt1, breaks = 30, probability = TRUE, main = “Monte Carlo Simulation Result of tt”,
    xlab = “simulated value”)
curve(dnorm(x, mean = mean(tt1), sd = sd(tt1)),
    col = “red”, lwd = 2, add = TRUE)
meanT = 4.194
sdT = 0.083
TT <- rnorm(n = 10000, mean = meanT, sd = sdT)
c(mean(TT), sd(TT))
t.test(tt1, TT)
~~~

## Acknowledgements

Patent pending (**63/984**,**445**), Dr. Philip P Foster, filed on 02/17/2026 with *USPTO/EPO*. This work was supported partly by a *National Science Foundation* Grant # *CBET* **2224942** to Dr. Aladin M Boriek.

## References

1 Rumelhart, D. E.; Hinton, G.E.; Williams, R.J. Learning representations by back-propagating errors. Nature 323, 3 (1986).

2 LeCun, Y., Bengio, Y. & Hinton, G. Deep learning. Nature 521, 436–444, doi:10.1038/nature14539 (2015).

3 Hinton, G. The Forward-Forward Algorithm: Some Preliminary Investigations arXiv, 16 (2022).

4 Hinton, G. E., Osindero, S. & Teh, Y. W. A fast learning algorithm for deep belief nets. Neural Comput 18, 1527–1554, doi:10.1162/neco.2006.18.7.1527 (2006).

5 Hinton, G. E. & Salakhutdinov, R. R. Reducing the dimensionality of data with neural networks. Science 313, 504–507, doi:10.1126/science.1127647 (2006).

6 Sánchez, V. et al. The Human Brain as a Combinatorial Complex. arXiv:2511.20692 (2025).< https://ui.adsabs.harvard.edu/abs/2025arXiv251120692S>.

7 Whittington, J. C. R. & Dorrell, W. How much neuroscience does a neuroscientist need to know?, arXiv:2601.02063 (2026). < https://ui.adsabs.harvard.edu/abs/2026arXiv260102063W>.

8 Rotenberg, V. S. Moravec’s paradox: consideration in the context of two brain hemisphere functions. Activitas Nervosa Superior 55.

9 Moravec, H. When will computer hardware match the human brain? Journal of Evolution and Technology 1 (1998).

10 Moravec, H. Robot. Mere Machine to Transcendent Mind. (Oxford University Press, 1998).

11 Noever, D. M. F. Moravec’s Paradox: Towards an Auditory Turing Test. arXiv arXiv:2507.23091v1 (2025).

12 Einstein, A. Zum gegenwärtigen Stande des Gravitationsproblems. Physikalische Zeitschrift 14, 1249–1262 (1913).

13 Einstein, A. Die Grundlage der allgemeinen Relativitätstheorie. Annalen der Physik 49, 769–822 (1916a).

14 Einstein, A. Näherungsweise Integration der Feldgleichungen der Gravitation. Königlich Preußische Akademie der Wissenschaften (Berlin). Sitzungsberichte, 688–696 (1916b).

15 Einstein A. Über Gravitationswellen. Königlich Preußische Akademie der Wissenschaften (Berlin). Sitzungsberichte, 154–167 (1918).

16 Foster, P. P., Conkin, J., Powell, M. R., Waligora, J. M. & Chhikara, R. S. Role of metabolic gases in bubble formation during hypobaric exposures. J.Appl.Physiol 84, 1088–1095 (1998).

17 Hosmer, D. W., Jr., Wang, C. Y., Lin, I. C. & Lemeshow, S. A computer program for stepwise logistic regression using maximum likelihood estimation. Comput Programs Biomed 8, 121–134, doi:10.1016/0010-468x(78)90047-8 (1978).

18 Lemeshow, S. & Hosmer, D. W., Jr. A review of goodness of fit statistics for use in the development of logistic regression models. Am J Epidemiol 115, 92–106, doi:10.1093/oxfordjournals.aje.a113284 (1982).

19 Hosmer, D. W., Hosmer, T., Le Cessie, S. & Lemeshow, S. A comparison of goodness-of-fit tests for the logistic regression model. Stat Med 16, 965–980, doi:10.1002/(sici)1097-0258(19970515)16:9<965::aid-sim509>3.0.co;2-o (1997).

20 Bengio, Y. L. P.; Popovici, D.; Larochelle, H. (Montreal, CA, 2006).

21 Hinton, G. E. D. L.; Yu, D.; Dahl, G.; Mohamed, A.; Jaitly, N.; Senior, A.; Vanhoucke, V.; Nguyen, P.; Sainath, T.; Kingsbury, B. in IEEE SIGNAL PROCESSING MAGAZINE 17 (IEEE, 2012).

22 Woodruff, K. K. et al. A pilot study for applying an extravehicular activity exercise prebreathe protocol to the International Space Station. NASA Technical Memorandum 210132 (2000).

23 M.L., G. et al. Design of a 2-hours prebreathe protocol for space walks from the International Space Sation. Aviat.Space Environ.Med. 71, 277 (2000).

24 Foster, P. P., Feiveson, A. H. & Boriek, A. M. Predicting time to decompression illness during exercise at altitude, based on formation and growth of bubbles. Am.J.Physiol Regul.Integr.Comp Physiol 279, R2317–R2328 (2000).

25 Foster, P. P., Feiveson, A. H., Glowinski, R., Izygon, M. & Boriek, A. M. A model for influence of exercise on formation and growth of tissue bubbles during altitude decompression. Am.J.Physiol Regul.Integr.Comp Physiol 279, R2304–R2316 (2000).

26 Foster, P. P. & Butler, B. D. Decompression to altitude: assumptions, experimental evidence, and future directions. J.Appl.Physiol 106, 678–690 (2009).

27 Glorot, X. B. A.; Bengio, Y. in Proceedings of the 14th International Conference on Artificial Intelligence and Statistics (AISTATS). (ed JMLR) (AISTATS).

28 Le Cun, Y. B. B.; Denker, J.S.; Henderson, D.; Howard, R.E.; Hubbard, W.; Jackel, L.D. in NIPS. 8 (NIPS).

29 Bona-Pellissier, J., Malgouyres, F. & Bachoc, F. Geometry-induced Regularization in Deep ReLU Neural Networks. arXiv:2402.08269 (2024). < https://ui.adsabs.harvard.edu/abs/2024arXiv240208269B>.

30 Grammer, J. K., Carrasco, M., Gehring, W. J. & Morrison, F. J. Age-related changes in error processing in young children: a school-based investigation. Dev Cogn Neurosci 9, 93–105, doi:10.1016/j.dcn.2014.02.001 (2014).

31 Bapu, A., Chen, T., Chien, C.-K. K., Muñoz Ewald, P. & Moore, A. G. Architecture independent generalization bounds for overparametrized deep ReLU networks. arXiv:2504.05695 (2025). < https://ui.adsabs.harvard.edu/abs/2025arXiv250405695B >.

32 Daniely, A. Deep Networks Learn Deep Hierarchical Models. arXiv:2601.00455 (2026). < https://ui.adsabs.harvard.edu/abs/2026arXiv260100455D >.

33 Simeone, O. Modern Neuromorphic AI: From Intra-Token to Inter-Token Processing. arXiv:2601.00245 (2026). < https://ui.adsabs.harvard.edu/abs/2026arXiv260100245S >.

34 Yu, Qian, Geisler, W. S. & Wei, X.-X. Quantifying task-relevant representational similarity using decision variable correlation. arXiv:2506.02164 (2025). < https://ui.adsabs.harvard.edu/abs/2025arXiv250602164Y>.

35 Alexander, S. in ACMS Conference Proceedings 2022. 10. (ed ACMS) 14 (ACMS, 2022).

36 Maier, M., Cheung, V. & Lieder, F. Learning from outcomes shapes reliance on moral rules versus cost-benefit reasoning. Nat Hum Behav, doi:10.1038/s41562-025-02271-w (2025).

37 Lake, B. M., Salakhutdinov, R. & Tenenbaum, J. B. Human-level concept learning through probabilistic program induction. Science 350, 1332–1338, doi:10.1126/science.aab3050 (2015).

38 Felleman, D. J. & Van Essen, D. C. Distributed hierarchical processing in the primate cerebral cortex. Cereb Cortex 1, 1–47, doi:10.1093/cercor/1.1.1-a (1991).

39 LeCun, Y. in A la Frontière de l’Intelligence Artificielle, des Sciences de la Connaissance et des Neurosciences. 599–604 (Cognitiva 85, Paris, 1985).

40 Freedman, D. J., Riesenhuber, M., Poggio, T. & Miller, E. K. Categorical representation of visual stimuli in the primate prefrontal cortex. Science 291, 312–316, doi:10.1126/science.291.5502.312 (2001).

41 Thorpe, S. J. & Fabre-Thorpe, M. Neuroscience. Seeking categories in the brain. Science 291, 260–263, doi:10.1126/science.1058249 (2001).

42 Cadieu, C. F. et al. Deep neural networks rival the representation of primate IT cortex for core visual object recognition. PLoS computational biology 10, e1003963, doi:10.1371/journal.pcbi.1003963 (2014).

43 Guerguiev, J., Lillicrap, T. P. & Richards, B. A. Towards deep learning with segregated dendrites. Elife 6, doi:10.7554/eLife.22901 (2017).

44 Richards, B. A. et al. A deep learning framework for neuroscience. Nature neuroscience 22, 1761–1770, doi:10.1038/s41593-019-0520-2 (2019).

45 Richards, B. A. & Lillicrap, T. P. Dendritic solutions to the credit assignment problem. Curr Opin Neurobiol 54, 28–36, doi:10.1016/j.conb.2018.08.003 (2019).

46 richardsRichards, B. A. et al. A deep learning framework for neuroscience. Nat Neurosci 22, 1761–1770, doi:10.1038/s41593-019-0520-2 (2019).

47 Lillicrap, T. P., Santoro, A., Marris, L., Akerman, C. J. & Hinton, G. Backpropagation and the brain. Nature reviews. Neuroscience 21, 335–346, doi:10.1038/s41583-020-0277-3 (2020).

48 Foster, P. P. et al. Protective mechanisms in hypobaric decompression. Aviation, space, and environmental medicine 84, 212–225 (2013).

49 Foster, P. P. in Right-to-left shunting, white matter hyperintensities: from altitude to diving. Patent foramen ovale & fitness to dive consensus (ed Richard E. Moon Alfred A. Bove) Ch. 3, 34–47 (UHMS DAN, 2016).

50 Tablante T.J. in How do Microchips work? How does your smartphone camera work? How does Bluetooth work? (ed Tablante T.J.) (Branch Education, 2021).

51 Kandala, A. et al. Hardware-efficient variational quantum eigensolver for small molecules and quantum magnets. Nature 549, 242–246, doi:10.1038/nature23879 (2017).

52 Voisin, B. et al. Valley interference and spin exchange at the atomic scale in silicon. Nature communications 11, 6124, doi:10.1038/s41467-020-19835-1 (2020).

53 Nakamura, Y., Pashkin, Y. A. & Tsai, J. S. Coherent control of macroscopic quantum states in a single-Cooper-pair box. Nature 398, 786–788, doi:10.1038/19718 (1999).

54 Devoret, M. H., Martinis, J. M., Esteve, D. & Clarke, J. Resonant Activation from the Zero-Voltage State of a Current-Biased Josephson Junction. Physical review letters 53, 1260–1263, doi:10.1103/PhysRevLett.53.1260 (1984).

55 Martinis, J. M., Devoret, M. H. & Clarke, J. Energy-Level Quantization in the Zero-Voltage State of a Current-Biased Josephson Junction. Physical review letters 55, 1543–1546, doi:10.1103/PhysRevLett.55.1543 (1985).

56 Devoret, M. H., Martinis, J. M. & Clarke, J. Measurements of Macroscopic Quantum Tunneling out of the Zero-Voltage State of a Current-Biased Josephson Junction. Physical review letters 55, 1908–1911, doi:10.1103/PhysRevLett.55.1908 (1985).

57 Melanson, D. et al. Thermodynamic computing system for AI applications. Nature communications 16, 3757, doi:10.1038/s41467-025-59011-x (2025).

58 He, Q. et al. Two-dimensional materials based two-transistor-two-resistor synaptic kernel for efficient neuromorphic computing. Nature communications 16, 4340, doi:10.1038/s41467-025-59815-x (2025).

59 Sebastian, A., Le Gallo, M., Khaddam-Aljameh, R. & Eleftheriou, E. Memory devices and applications for in-memory computing. Nat Nanotechnol 15, 529–544, doi:10.1038/s41565-020-0655-z (2020).

60 Wang, X.-D. et al. Multiscale simulations of growth-dominated Sb2Te phase-change material for non-volatile photonic applications. npj Computational Materials 9, 136, doi:10.1038/s41524-023-01098-1 (2023).

61 Yu, J. et al. Simultaneously ultrafast and robust two-dimensional flash memory devices based on phase-engineered edge contacts. Nature communications 14, 5662, doi:10.1038/s41467-023-41363-x (2023).

62 Park, M. K. et al. Charge-trap synaptic device with polycrystalline silicon channel for low power in-memory computing. Scientific reports 14, 29089, doi:10.1038/s41598-024-80272-x (2024).

63 Harris, J. J., Jolivet, R., Engl, E. & Attwell, D. Energy-Efficient Information Transfer by Visual Pathway Synapses. Curr Biol 25, 3151–3160, doi:10.1016/j.cub.2015.10.063 (2015).

64 Hodgkin, A. L. & Huxley, A. F. A quantitative description of membrane current and its application to conduction and excitation in nerve. The Journal of physiology 117, 500–544, doi:10.1113/jphysiol.1952.sp004764 (1952).

65 Schmidt, H. & Knosche, T. R. Action potential propagation and synchronisation in myelinated axons. PLoS computational biology 15, e1007004, doi:10.1371/journal.pcbi.1007004 (2019).

66 Hu, H. W. M. Action Potential Modulation of Neural Spin Networks Suggests Possible Role of Spin. CERN (European Council for Nuclear Research) - EXT-2004-015, 10 (2004).

67 Hu, H. W. M. Possible Roles of Neural Electron Spin Networks in Memory and Consciousness. CERN (European Council for Nuclear Research) - EXT-2004-027, 19 (2004).

68 Yang, R., Ping, H., Xiao, X., Kiani, R. & Bogdan, P. Spiking dynamics of individual neurons reflect changes in the structure and function of neuronal networks. Nature communications 16, 6994, doi:10.1038/s41467-025-62202-1 (2025).

69 Pazos, S. et al. Synaptic and neural behaviours in a standard silicon transistor. Nature 640, 69–76, doi:10.1038/s41586-025-08742-4 (2025).

70 Hinton, G. E., Dayan, P., Frey, B. J. & Neal, R. M. The “wake-sleep” algorithm for unsupervised neural networks. Science 268, 1158–1161, doi:10.1126/science.7761831 (1995).

71 Herculano-Houzel, S. The remarkable, yet not extraordinary, human brain as a scaled-up primate brain and its associated cost. Proceedings of the National Academy of Sciences of the United States of America 109 Suppl 1, 10661–10668, doi:10.1073/pnas.1201895109 (2012).

72 Eslami, S. M. A. et al. Neural scene representation and rendering. Science 360, 1204–1210, doi:10.1126/science.aar6170 (2018).

73 Mnih, V. et al. Human-level control through deep reinforcement learning. Nature 518, 529–533, doi:10.1038/nature14236 (2015).

74 Tenenbaum, J. B., Kemp, C., Griffiths, T. L. & Goodman, N. D. How to grow a mind: statistics, structure, and abstraction. Science 331, 1279–1285, doi:10.1126/science.1192788 (2011).

75 Le Bihan, D. Looking into the functional architecture of the brain with diffusion MRI. Nature reviews. Neuroscience 4, 469–480, doi:10.1038/nrn1119 (2003).

76 Taylor, H. P. et al. Functional hierarchy of the human neocortex across the lifespan. Nature, doi:10.1038/s41586-026-10219-x (2026).

77 Kringelbach, M. L. et al. Dynamic coupling of whole-brain neuronal and neurotransmitter systems. Proceedings of the National Academy of Sciences of the United States of America 117, 9566–9576, doi:10.1073/pnas.1921475117 (2020).

78 Kandala, A. et al. Error mitigation extends the computational reach of a noisy quantum processor. Nature 567, 491–495, doi:10.1038/s41586-019-1040-7 (2019).

79 Havlicek, V. et al. Supervised learning with quantum-enhanced feature spaces. Nature 567, 209–212, doi:10.1038/s41586-019-0980-2 (2019).

80 Scott, R. S. et al. Neuronal adaptation involves rapid expansion of the action potential initiation site. Nature communications 5, 3817, doi:10.1038/ncomms4817 (2014).

81 Ford, M. C. et al. Tuning of Ranvier node and internode properties in myelinated axons to adjust action potential timing. Nature communications 6, 8073, doi:10.1038/ncomms9073 (2015).

82 El Hady, A. & Machta, B. B. Mechanical surface waves accompany action potential propagation. Nature communications 6, 6697, doi:10.1038/ncomms7697 (2015).

83 Bernaerts, Y. et al. Combined statistical-biophysical modeling links ion channel genes to physiology of cortical neuron types. bioRxiv, doi:10.1101/2023.03.02.530774 (2025).

84 Hafting, T., Fyhn, M., Molden, S., Moser, M. B. & Moser, E. I. Microstructure of a spatial map in the entorhinal cortex. Nature 436, 801–806, doi:10.1038/nature03721 (2005).

85 Foster, P. P. Role of physical and mental training in brain network configuration. Frontiers in aging neuroscience 7, 117, doi:10.3389/fnagi.2015.00117 (2015).

86 Jezek, K., Henriksen, E. J., Treves, A., Moser, E. I. & Moser, M. B. Theta-paced flickering between place-cell maps in the hippocampus. Nature 478, 246–249 (2011).

87 Colgin, L. L. et al. Attractor-map versus autoassociation based attractor dynamics in the hippocampal network. J.Neurophysiol. 104, 35–50 (2010).

88 Ito, H. T., Zhang, S. J., Witter, M. P., Moser, E. I. & Moser, M. B. A prefrontal-thalamo-hippocampal circuit for goal-directed spatial navigation. Nature 522, 50–55, doi:10.1038/nature14396 (2015).

89 Morris, R. G., Garrud, P., Rawlins, J. N. & O’Keefe, J. Place navigation impaired in rats with hippocampal lesions. Nature 297, 681–683, doi:10.1038/297681a0 (1982).

90 O’Keefe, J. & Burgess, N. Geometric determinants of the place fields of hippocampal neurons. 35. Nature 381, 425–428 (1996).

91 O’Keefe, J. & Burgess, N. Geometric determinants of the place fields of hippocampal neurons. Nature 381, 425–428, doi:10.1038/381425a0 (1996).

92 Thorpe, S., Fize, D. & Marlot, C. Speed of processing in the human visual system. Nature 381, 520–522, doi:10.1038/381520a0 (1996).

93 Bharadwaj, R. et al. Conserved higher-order chromatin regulates NMDA receptor gene expression and cognition. Neuron 84, 997–1008, doi:10.1016/j.neuron.2014.10.032 (2014).

94 Einstein, A. P. B.; Rosen, N. Can quantum-mechanical description of physical reality be considered complete? Phys. Rev. 47 (1935).

95 Rao, R. P. & Sejnowski, T. J. Complex Cell-like Direction Selectivity through Spike-Timing Dependent Plasticity. IETE J Res 49, 97–111, doi:10.1080/03772063.2003.11416329 (2003).

96 Sejnowski, T. J. Thinking about thinking: AI offers theoretical insights into human memory. The Transmitter, 14 (2025).

97 Lillicrap, T. P., Cownden, D., Tweed, D. B. & Akerman, C. J. Random synaptic feedback weights support error backpropagation for deep learning. Nature communications 7, 13276, doi:10.1038/ncomms13276 (2016).

98 Lake, B. M. & Baroni, M. Human-like systematic generalization through a meta-learning neural network. Nature 623, 115–121, doi:10.1038/s41586-023-06668-3 (2023).

99 Park, H. J. & Friston, K. Structural and functional brain networks: from connections to cognition. Science 342, 1238411, doi:10.1126/science.1238411 (2013).

100 Brobey, R. K. et al. Klotho Protects Dopaminergic Neuron Oxidant-Induced Degeneration by Modulating ASK1 and p38 MAPK Signaling Pathways. PloS one 10, e0139914, doi:10.1371/journal.pone.0139914 (2015).

101 Pardo, P. S. & Boriek, A. M. SIRT1 Regulation in Ageing and Obesity. Mech Ageing Dev 188, 111249, doi:10.1016/j.mad.2020.111249 (2020).

102 Brobey, R. K., Dheghani, M., Foster, P. P., Kuro-o, M. & Rosenblatt, K. P. Klotho Regulates 14-3-3ζ Monomerization and Binding to the ASK1 Signaling Complex in Response to Oxidative Stress. PloS one 10, e0141968, doi:10.1371/journal.pone.0141968 (2015).

103 Dobbin, M. M. et al. SIRT1 collaborates with ATM and HDAC1 to maintain genomic stability in neurons. Nature neuroscience 16, 1008–1015, doi:10.1038/nn.3460 (2013).

104 Baker, C. L. J.-E. J.; Saxe, R.; Tenenbaum, J.B. Rational quantitative attribution of beliefs, desires and percepts in human mentalizing. Nature human behaviour 01, 10, doi:10.1038/s41562-017-0064 (2017).

105 Buchel, C., Coull, J. T. & Friston, K. J. The predictive value of changes in effective connectivity for human learning. Science 283, 1538–1541, doi:10.1126/science.283.5407.1538 (1999).

106 Heeger, D. J. & Ress, D. What does fMRI tell us about neuronal activity? Nature reviews. Neuroscience 3, 142–151, doi:10.1038/nrn730 (2002).

107 Logothetis, N. K. The neural basis of the blood-oxygen-level-dependent functional magnetic resonance imaging signal. Philos.Trans.R.Soc.Lond B Biol.Sci. 357, 1003–1037 (2002).

108 Logothetis, N. K. What we can do and what we cannot do with fMRI. Nature 453, 869–878 (2008).

109 Kandel, E. R. The molecular biology of memory storage: a dialogue between genes and synapses. Science 294, 1030–1038, doi:10.1126/science.1067020 (2001).

110 Kandel, E. R. The biology of memory: a forty-year perspective. The Journal of neuroscience : the official journal of the Society for Neuroscience 29, 12748–12756, doi:10.1523/JNEUROSCI.3958-09.2009 (2009).

111 Zheng, J. & Meister, M. The unbearable slowness of being: Why do we live at 10 bits/s? Neuron 113, 192–204, doi:10.1016/j.neuron.2024.11.008 (2025).

112 Chaudhuri, R. & Fiete, I. Computational principles of memory. Nature neuroscience 19, 394–403, doi:10.1038/nn.4237 (2016).

113 Kim, J. H., Daie, K. & Li, N. A combinatorial neural code for long-term motor memory. Nature 637, 663–672, doi:10.1038/s41586-024-08193-3 (2025).

114 Suzuki, A. et al. A cortical cell ensemble in the posterior parietal cortex controls past experience-dependent memory updating. Nature communications 13, 41, doi:10.1038/s41467-021-27763-x (2022).

115 Jones, S. G. et al. A memory transcriptome time course reveals essential long-term memory transcription factors. Nature communications 16, 9320, doi:10.1038/s41467-025-64379-x (2025).

116 Wu, Y. & Maass, W. A simple model for Behavioral Time Scale Synaptic Plasticity (BTSP) provides content addressable memory with binary synapses and one-shot learning. Nature communications 16, 342, doi:10.1038/s41467-024-55563-6 (2025).

117 Theparambil, S. M. et al. Adenosine signalling to astrocytes coordinates brain metabolism and function. Nature 632, 139–146, doi:10.1038/s41586-024-07611-w (2024).

118 Tome, D. F. et al. Dynamic and selective engrams emerge with memory consolidation. Nature neuroscience 27, 561–572, doi:10.1038/s41593-023-01551-w (2024).

119 Hadzibegovic, S. et al. Early intrinsic excitability plasticity of neocortical engram neurons defines memory formation and precision. Nature communications, doi:10.1038/s41467-025-66975-3 (2025).

120 Li, Z. et al. Locating causal hubs of memory consolidation in spontaneous brain network in male mice. Nature communications 14, 5399, doi:10.1038/s41467-023-41024-z (2023).

121 de Ceglia, R. et al. Specialized astrocytes mediate glutamatergic gliotransmission in the CNS. Nature 622, 120–129, doi:10.1038/s41586-023-06502-w (2023).

122 Jeong, Y. et al. Synaptic plasticity-dependent competition rule influences memory formation. Nature communications 12, 3915, doi:10.1038/s41467-021-24269-4 (2021).

123 Colgin, L. L. & Moser, E. I. Neuroscience: rewinding the memory record. Nature 440, 615–617 (2006).

124 Derdikman, D. et al. Fragmentation of grid cell maps in a multicompartment environment. Nature neuroscience 12, 1325–1332, doi:10.1038/nn.2396 (2009).

125 Bonnevie, T. et al. Grid cells require excitatory drive from the hippocampus. Nature neuroscience 16, 309–317, doi:10.1038/nn.3311 (2013).

126 Pedamonti, D. et al. Hippocampus supports multi-task reinforcement learning under partial observability. Nature communications 16, 9619, doi:10.1038/s41467-025-64591-9 (2025).

127 Busch, A. et al. Neuronal activation sequences in lateral prefrontal cortex encode visuospatial working memory during virtual navigation. Nature communications 15, 4471, doi:10.1038/s41467-024-48664-9 (2024).

128 Marco, A. et al. Mapping the epigenomic and transcriptomic interplay during memory formation and recall in the hippocampal engram ensemble. Nature neuroscience 23, 1606–1617, doi:10.1038/s41593-020-00717-0 (2020).

129 Stott, R. T., Kritsky, O. & Tsai, L. H. Profiling DNA break sites and transcriptional changes in response to contextual fear learning. PloS one 16, e0249691, doi:10.1371/journal.pone.0249691 (2021).

130 Jovasevic, V. et al. Formation of memory assemblies through the DNA-sensing TLR9 pathway. Nature 628, 145–153, doi:10.1038/s41586-024-07220-7 (2024).

131 Herber, C. S., Pratt, K. J. B., Shea, J. M., Villeda, S. A. & Giocomo, L. M. Spatial coding dysfunction and network instability in the aging medial entorhinal cortex. Nature communications 16, 8770, doi:10.1038/s41467-025-63229-0 (2025).

132 Xu, Z., Geron, E., Perez-Cuesta, L. M., Bai, Y. & Gan, W. B. Generalized extinction of fear memory depends on co-allocation of synaptic plasticity in dendrites. Nature communications 14, 503, doi:10.1038/s41467-023-35805-9 (2023).

133 Zaki, Y. et al. Offline ensemble co-reactivation links memories across days. Nature 637, 145–155, doi:10.1038/s41586-024-08168-4 (2025).

134 Yount, S. T. et al. Parallel neuronal structural plasticity with memory trace formation in the orbitofrontal cortex. Nature communications 16, 8521, doi:10.1038/s41467-025-63542-8 (2025).

135 Park, G. et al. Hippocampal-cortical interactions in the consolidation of social memory. Nature communications 16, 8430, doi:10.1038/s41467-025-64264-7 (2025).

136 Higgs, P. W. Broken Symmetries and the Masses of Gauge Bosons. Physical review letters 13, 508–509, doi:10.1103/PhysRevLett.13.508 (1964).

137 Higgs, P. W. Spontaneous Symmetry Breakdown without Massless Bosons. Physical Review 145, 1156–1163, doi:10.1103/PhysRev.145.1156 (1966).

138 Looser, Z. J. et al. Oligodendrocyte-axon metabolic coupling is mediated by extracellular K(+) and maintains axonal health. Nature neuroscience 27, 433–448, doi:10.1038/s41593-023-01558-3 (2024).

139 McNamara, N. B. et al. Microglia regulate central nervous system myelin growth and integrity. Nature 613, 120–129, doi:10.1038/s41586-022-05534-y (2023).

140 Hua, H., Gu, H., Ma, K., Jia, Y. & Wu, L. Dynamics and conditions for inhibitory synaptic current to induce bursting and spreading depolarization in pyramidal neurons. Scientific reports 15, 8886, doi:10.1038/s41598-025-92647-9 (2025).

141 Nielsen, B. S. et al. Glial Versus Neuronal Na(+)/K(+) -ATPase in Activity-Evoked K(+) Clearance and Their Sensitivity to Elevated Extracellular K(). Glia 73, 1805–1816, doi:10.1002/glia.70034 (2025).

142 Steinmetz, N. A. et al. Neuropixels 2.0: A miniaturized high-density probe for stable, long-term brain recordings. Science 372, doi:10.1126/science.abf4588 (2021).

143 Ye, Z. et al. Ultra-high-density Neuropixels probes improve detection and identification in neuronal recordings. Neuron 113, 3966–3982 e3912, doi:10.1016/j.neuron.2025.08.030 (2025).

144 Penrose, R. L. M.S.; Shimony, A.; Cartwright Hawking, S. The Large, the Small and the Human Mind. First Edition edn, (Cambridge University Press, 1997).

145 Penrose, R. H. S. Consciousness in the Universe: Neuroscience, Quantum Space-Time Geometry and Orch OR Theory. Journal of Cosmology 14 (2011).

146 Hameroff, S. & Penrose, R. Consciousness in the universe: a review of the ‘Orch OR’ theory. Phys Life Rev 11, 39–78, doi:10.1016/j.plrev.2013.08.002 (2014).

147 Stapp, H. P. Quantum propensities and the brain-mind connection. Foundations of Physics 21, 1451–1477, doi:10.1007/BF01889652 (1991).

148 Zeh, H. D. The Problem of Conscious Observation in Quantum Mechanical Description. quant-ph/9908084 (1999). < https://ui.adsabs.harvard.edu/abs/1999quant.ph.8084Z >.

149 Zurek, W. H. Decoherence and the transition from quantum to classical --REVISITED. quant-ph/0306072 (2003). < https://ui.adsabs.harvard.edu/abs/2003quant.ph.6072Z >.

150 Tegmark, M. Importance of quantum decoherence in brain processes. Physical Review E 61, 4194–4206, doi:10.1103/PhysRevE.61.4194 (2000).

151 Yamauchi, A. et al. Room-temperature quantum coherence of entangled multiexcitons in a metal-organic framework. Science Advances 10, eadi3147, doi:10.1126/sciadv.adi3147 (2024).

152 Arancibia-Carcamo, I. L. & Attwell, D. The node of Ranvier in CNS pathology. Acta neuropathologica 128, 161–175, doi:10.1007/s00401-014-1305-z (2014).

153 Poliak, S. & Peles, E. The local differentiation of myelinated axons at nodes of Ranvier. Nature Reviews Neuroscience 4, 968–980, doi:10.1038/nrn1253 (2003).

154 Rasband, M. N. & Peles, E. Mechanisms of node of Ranvier assembly. Nature Reviews Neuroscience 22, 7–20, doi:10.1038/s41583-020-00406-8 (2021).

155 Lubetzki, C., Sol-Foulon, N. & Desmazières, A. Nodes of Ranvier during development and repair in the CNS. Nature Reviews Neurology 16, 426–439, doi:10.1038/s41582-020-0375-x (2020).

156 Morán, O. & Mateu, L. Loosening of paranodal myelin by repetitive propagation of action potentials. Nature 304, 344–345, doi:10.1038/304344a0 (1983).

157 Osso, L. A. & Hughes, E. G. Dynamics of mature myelin. Nature neuroscience 27, 1449–1461, doi:10.1038/s41593-024-01642-2 (2024).

158 Meschkat, M. et al. White matter integrity in mice requires continuous myelin synthesis at the inner tongue. Nature communications 13, 1163, doi:10.1038/s41467-022-28720-y (2022).

159 Djannatian, M. et al. Two adhesive systems cooperatively regulate axon ensheathment and myelin growth in the CNS. Nature communications 10, 4794, doi:10.1038/s41467-019-12789-z (2019).

160 Scherer, S. S. Nodes, paranodes, and incisures: from form to function. Ann N Y Acad Sci 883, 131–142 (1999).

161 Scherer, S. S. & Arroyo, E. J. Recent progress on the molecular organization of myelinated axons. Journal of the Peripheral Nervous System 7, 1–12, doi:10.1046/j.1529-8027.2002.02001.x (2002).

162 Elbaz, B. et al. The bone transcription factor Osterix controls extracellular matrix- and node of Ranvier-related gene expression in oligodendrocytes. Neuron 112, 247–263.e246, doi:10.1016/j.neuron.2023.10.008 (2024).

163 Devaux, J. J. & Faivre-Sarrailh, C. Neuro-glial interactions at the nodes of Ranvier: implication in health and diseases. Frontiers in cellular neuroscience Volume 7 - 2013 (2013).

164 Cullen, C. L. et al. Periaxonal and nodal plasticities modulate action potential conduction in the adult mouse brain. Cell reports 34, 108641, doi:10.1016/j.celrep.2020.108641 (2021).

165 Shimizu, T. et al. Oligodendrocyte dynamics dictate cognitive performance outcomes of working memory training in mice. Nature communications 14, 6499, doi:10.1038/s41467-023-42293-4 (2023).

166 Markram, H., Lubke, J., Frotscher, M., Roth, A. & Sakmann, B. Physiology and anatomy of synaptic connections between thick tufted pyramidal neurones in the developing rat neocortex. The Journal of physiology 500 ( Pt 2), 409–440, doi:10.1113/jphysiol.1997.sp022031 (1997).

167 Arcas, B. A. y., Fairhall, A. L. & Bialek, W. Computation in a Single Neuron: Hodgkin and Huxley Revisited. Neural Computation 15, 1715–1749, doi:10.1162/08997660360675017 (2003).

168 Sachdev, S. Quantum Phase Transitions. 2 edn, (Cambridge University Press, 2011).

169 Heyl, M. Dynamical quantum phase transitions: a review. Reports on Progress in Physics 81, 054001, doi:10.1088/1361-6633/aaaf9a (2018).

170 Lee, M.-W. & Hsu, L.-Y. Polariton-assisted resonance energy transfer beyond resonant dipole-dipole interaction: A transition-current-density approach. Physical Review A 107, 053709, doi:10.1103/PhysRevA.107.053709 (2023).

171 Okamura, Y. et al. Photovoltaic effect by soft phonon excitation. Proceedings of the National Academy of Sciences 119, e2122313119, doi:10.1073/pnas.2122313119 (2022).

172 Xu, C. & Zong, A. Time-domain study of coupled collective excitations in quantum materials. npj Quantum Materials 10, 21, doi:10.1038/s41535-025-00726-x (2025).

173 Ng, R. C. et al. Excitation and detection of acoustic phonons in nanoscale systems. Nanoscale 14, 13428–13451, doi:10.1039/D2NR04100F (2022).

174 Panagopoulos, D. J., Johansson, O. & Carlo, G. L. Polarization: A Key Difference between Man-made and Natural Electromagnetic Fields, in regard to Biological Activity. Scientific reports 5, 14914, doi:10.1038/srep14914 (2015).

175 Project, C. et al. Single-Electron Detection and Spectroscopy via Relativistic Cyclotron Radiation. Physical review letters 114, 162501, doi:10.1103/PhysRevLett.114.162501 (2015).

176 Zhang, Y. et al. Ultrafast and hypersensitive phase imaging of propagating internodal current flows in myelinated axons and electromagnetic pulses in dielectrics. Nature communications 13, 5247, doi:10.1038/s41467-022-33002-8 (2022).

177 Cai, Y. et al. Myelin–axon interface vulnerability in Alzheimer’s disease revealed by subcellular proteomics and imaging of human and mouse brain. Nature neuroscience 28, 1418–1435, doi:10.1038/s41593-025-01973-8 (2025).

178 Pravata, E. et al. Biometry extraction and probabilistic anatomical atlas of the anterior Visual Pathway using dedicated high-resolution 3-D MRI. Scientific reports 14, 453, doi:10.1038/s41598-023-50980-x (2024).

179 Takemura, H. et al. Occipital White Matter Tracts in Human and Macaque. Cereb Cortex 27, 3346–3359, doi:10.1093/cercor/bhx070 (2017).

180 West, K. L. et al. BOLD hemodynamic response function changes significantly with healthy aging. NeuroImage 188, 198–207, doi:10.1016/j.neuroimage.2018.12.012 (2019).

181 Le Bihan, D. From Brownian motion to virtual biopsy: a historical perspective from 40 years of diffusion MRI. Jpn J Radiol 42, 1357–1371, doi:10.1007/s11604-024-01642-z (2024).

182 Peltier, J., Travers, N., Destrieux, C. & Velut, S. Optic radiations: a microsurgical anatomical study. Journal of neurosurgery 105, 294–300, doi:10.3171/jns.2006.105.2.294 (2006).

183 Sawilowsky, S. S. “Fermat, Schubert, Einstein, and Behrens-Fisher: The Probable Difference Between Two Means With Different Variances”. J. Modern Applied Statistical Methods 1, 11 (2002).

184 Roussel, T., Frydman, L., Le Bihan, D. & Ciobanu, L. Brain sugar consumption during neuronal activation detected by CEST functional MRI at ultra-high magnetic fields. Scientific reports 9, 4423, doi:10.1038/s41598-019-40986-9 (2019).

185 Boido, D. et al. Mesoscopic and microscopic imaging of sensory responses in the same animal. Nature communications 10, 1110, doi:10.1038/s41467-019-09082-4 (2019).

186 Abe, Y., Tsurugizawa, T., Le Bihan, D. & Ciobanu, L. Spatial contribution of hippocampal BOLD activation in high-resolution fMRI. Scientific reports 9, 3152, doi:10.1038/s41598-019-39614-3 (2019).

187 de Sousa, A. F. et al. The prefrontal cortex controls memory organization in the hippocampus. Nature neuroscience, doi:10.1038/s41593-026-02231-1 (2026).

