## Supplementary figures and images for "Charge-trap flash memory cells of the brain"

### Fig9

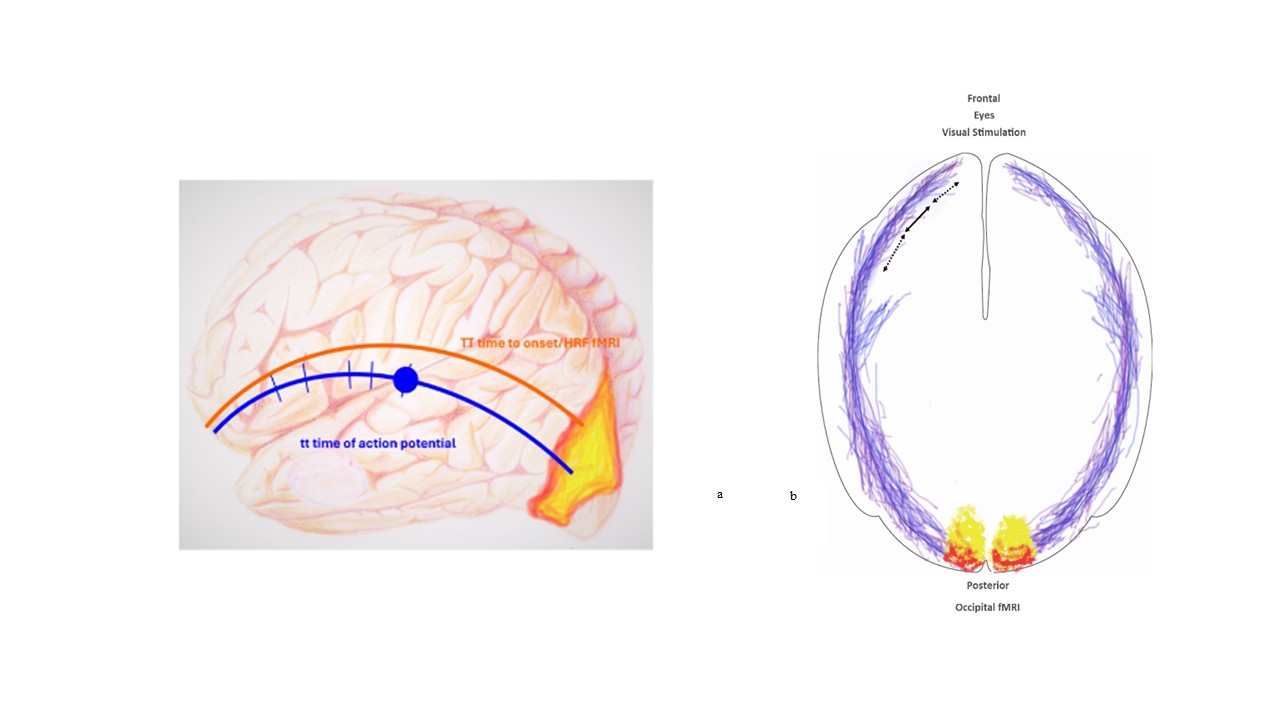
